# From Maps to Models: A Survey on the Reliability of Small Studies of Task-Based fMRI

**DOI:** 10.1101/2024.08.05.606611

**Authors:** Patrick Sadil, Martin A. Lindquist

## Abstract

Task-based functional magnetic resonance imaging (fMRI) is a powerful tool for studying brain function. However, the reliability and viability of small-sample studies remain a concern. While it is well understood that larger samples are preferable, researchers often need to interpret findings from small studies (e.g., when reviewing the literature, analyzing pilot data, or assessing subsamples). However, quantitative guidance for making these judgments remains scarce. To address this gap, we leverage the Human Connectome Project’s Young Adult and UK Biobank datasets to survey a range of standard task-based fMRI analyses, from obtaining regional activation maps to performing predictive modeling. These analyses are repeated using volumetric and two types of cortical surface data. For classic mass-univariate analyses (e.g., regional activation detection or cluster peak localization), studies with as few as 40 participants can be adequate depending on the effect size. For predictive modeling, similar sample sizes can be used to detect whether a feature is predictable, but developing stable, generalizable models typically requires cohorts at least an order of magnitude larger, and possibly two (hundreds or thousands). Together, these results clarify how reliability depends on the interplay of effect size, sample size, and analysis type, offering practical guidance for designing and interpreting small-scale task-fMRI studies.

## 1 Introduction

Considerable attention has focused on problems caused by inadequate sample sizes in task-based functional magnetic resonance imaging (fMRI) studies (e.g., Bossier et al., 2020; Button et al., 2013; M. A. Lindquist et al., 2013; Lohmann et al., 2017; Marek et al., 2022; Poldrack et al., 2017; Thirion et al., 2007; Turner et al., 2018; Yarkoni, 2009). Studies with too few participants tend to produce noisy effect estimates, which can inflate both false-positive and false-negative rates (Cremers et al., 2017; Gonzalez-Castillo et al., 2012; Lieberman & Cunningham, 2009). Furthermore, effect sizes estimated from significant results are often exaggerated, and the replicability of activation and significance maps is low (Bossier et al., 2020; Marek et al., 2022; Reddan et al., 2017; Turner et al., 2018; Yarkoni, 2009). Together, these issues can impede cumulative scientific progress (see also Ottenbacher, 1996; Schmidt, 1996).

In response, the neuroimaging community has developed a range of solutions. These include advanced methodological solutions (e.g., meta-analytic techniques; Eickhoff et al., 2009; Turkeltaub et al., 2002; Wager et al., 2003) and social improvements (e.g., large-scale data collections with hundreds to tens of thousands of participants; Di Martino et al., 2014; Krieger et al., 2017; Miller et al., 2016; Van Essen et al., 2013; Volkow et al., 2018). While these efforts promise a future of more valid and reliable research, the need to interpret small studies persists, as researchers must continue to rely on older publications, analyze pilot data, and work with limited clinical populations. Given the prevalence of small studies in the neuroimaging literature (Poldrack et al., 2017), it is crucial to have well-calibrated expectations for what they can and cannot tell us.

Notably, small task-based studies can still support robust research in several specific ways (Bossier et al., 2020; Kragel et al., 2021; Woo & Wager, 2016). For example, multivariate models trained using only tens of participants can exhibit strong out-of-sample performance on external cohorts of hundreds of participants (Han et al., 2022; J.-J. Lee et al., 2021; Wager et al., 2013; Woo & Wager, 2016). When multivariate models are tested appropriately – such as through cross-validation and, ideally, external test sets – predictive performance in smaller datasets can justify further investigation; many research questions can be answered based on whether, and not how well, fMRI data support predictions (Naselaris et al., 2011). That said, larger and more diverse training generally improves performance (Chen et al., 2023; Greene et al., 2022; He et al., 2020; Schulz et al., 2022; Traut et al., 2022), and small samples usually impede accurate estimation of generalization performance (Poldrack et al., 2020; Varoquaux, 2017).

Classic mass-univariate approaches can also yield high-quality results from small samples. For example, high power can be achieved by focusing on robust effects (Desmond & Glover, 2002; J. N. Lee et al., 2010), which are prevalent in regions like the somatomotor or visual networks (Engel et al., 1994; Grodd et al., 2001). With optimized designs, single-subject run-to-run reliability is sufficiently high to support clinical applications, such as surgical mapping (Brannen et al., 2001; Fernandez et al., 2003). Methodological advances are also improving what can be extracted from smaller datasets. While traditional voxel-wise, regional, or cluster-based inference replicates poorly in small samples (Nee, 2019; Turner et al., 2018, 2019), several alternative methods can improve both sensitivity and specificity (Noble et al., 2020; Spisák et al., 2019; Wang et al., 2021). As with multivariate methods, this success suggests that a blanket dismissal of small studies is unwarranted and that even modest sample sizes can support rigorous science.

In this paper, we survey the validity and reliability of inferences from task-based fMRI conducted on small samples (that is, fewer than 100 participants). We evaluate four analysis levels spanning statistical maps to predictive models. First, we considered region-of-interest (ROI) analyses using atlas-based parcellations. Second, we considered peak activity localization. Third, we explored topographic analyses of pointwise (voxel or vertex) effect size maps (Misic, 2025). Fourth, we assessed analyses based on multivariate modeling.

For each analysis level, we used a three-step procedure (Bossier et al., 2020; Cremers et al., 2017; Geuter et al., 2018; Lohmann et al., 2017; Thirion et al., 2007; Turner et al., 2018). First, we built and described a “gold standard” – a population about which individual studies should support inference. Second, we generated pseudo-studies via bootstrapping from the gold standard across a range of sample sizes and assessed validity by comparing with the gold standard. Third, we evaluated the reliability of the studies by measuring consistency across resamples, using, in most cases, the intra-class correlation coefficient (Noble et al., 2021).

## 2 Methods

Analyses relied on two large-scale datasets: the Human Connectome Project Young Adult cohort (HCP-YA; Feinberg et al., 2010; Moeller et al., 2010; Setsompop et al., 2012; Van Essen et al., 2013; Xu et al., 2012) and the UK Biobank (UKB; Alfaro-Almagro et al., 2018). Below, we summarize these datasets and outline the corresponding analysis pipelines. The tasks and contrasts included in analyses are described in the Supplementary Materials (subsubsection 5.2.1).

### 2.1 Data

Analyses used the HCP Young Adult and UK Biobank datasets, described in the following sections. For an overview of the participants, see Table 1.

**Table 1:** Participant Characteristics. ^*1*^ Median (Q1, Q3); n (%)

|  | HCP-YA | UKB |
| --- | --- | --- |
| Characteristic | N = 384<sup>1</sup> | N = 42,578<sup>1</sup> |
| age | 29 (26, 32) | 64 (58, 70) |
| sex |  |  |
| Female | 216 (56%) | 22,590 (53%) |
| Male | 168 (44%) | 19,988 (47%) |

#### 2.1.1 HCP 500

The Human Connectome Project 500 (HCP 500) consists of both structural and functional data from approximately 500 participants. Although the full HCP-YA dataset comprises over 1200 participants, the 500-participant release was used because our survey requires the results of the volumetric pipelines that HCP only provides for the S500 release.

Because the HCP-YA dataset includes sibling pairs (including twins) that may impact the generalizability of our analyses, we retained only one individual per twin pair. Additionally, any scans flagged by the HCP-YA QC procedure were excluded. For a list of excluded participants, see the Supplementary Materials (subsection 5.1). In the resulting subset (384 participants), average framewise displacement was 0.17 mm (SD: 0.0739). No volumes were scrubbed.

All data were acquired on a Siemens Skyra 3T scanner at Washington University in St. Louis. For each task, two runs were acquired: one with a right-to-left phase encoding and the other with a left-to-right phase encoding. Whole-brain echo-planar imaging acquisitions were acquired with a 32-channel head coil with TR=720 ms, TE=33.1 ms, flip angle=52^*°*^, bandwidth=2290 Hz*/*Px, in-place field-of-view=208*×*180 mm, 72 slices, 2 mm isotropic voxels, with a multi-band acceleration factor of 8. For a complete description of the fMRI data acquisition, see Van Essen et al. (2013).

Scans were preprocessed according to the HCP “fMRIVolume” minimal-preprocessing pipeline (Glasser et al., 2013), which includes gradient unwarping, motion correction, fieldmap-based distortion correction, brain-boundary-based registration to a structural T1-weighted scan, non-linear registration into MNI152 space, grand-mean intensity normalization, and spatial smoothing using a Gaussian kernel with a full-width half-maximum of 4 mm. Analyses were restricted to either a whole-brain mask or a graymatter mask that included both cortical and subcortical voxels.

Several analyses were performed on contrast maps from a general linear model provided by the HCP (Barch et al., 2013). For each task, predictors (described for each task in subsubsection 5.2.1) were convolved with a canonical hemodynamic response function to generate regressors. To compensate for slice-timing differences and variability in the hemodynamic delay across regions, temporal derivatives were included and treated as variables of no interest. Both the data and the design matrix were temporally filtered using a linear high-pass filter (cutoff 200 s). During model fitting, the time series was pre-whitened. For each task, we analyzed a single contrast of parameter estimates (the result of a fixed-effects analysis on run-wise “Level 1” analyses). Although this approach did not include denoising strategies standard in many analysis pipelines (e.g., including motion regressors as confounds), prior work finds that denoising these data does not substantially improve individual-level *z*-statistics (Barch et al., 2013).

For each contrast map, we analyzed the volumetric data (VOL), surface data that had been registered using traditional methods (SURFACE), and surface data that had been registered using multimodal surface matching (MSMAll; Robinson et al., 2014, 2018).

#### 2.1.2 UKB

The UK Biobank (UKB) also comprises both structural and functional data. The data were downloaded in January 2024 and consisted of more than 40,000 participants with usable data (participants with a value for Field 25733: “Amount of warping applied to non-linearly align T1 brain image to standard-space”).

For additional details on the acquisition and preprocessing, see the report by Alfaro-Almagro et al. (2018). Information about the task is available in subsubsection 5.2.2. Average framewise displacement was 0.25 mm (SD: 0.115). No volumes were scrubbed. UKB “Cognitive” variables were extracted using the FMRIB UKBiobank Normalisation, Parsing And Cleaning Kit (McCarthy, 2023).

### 2.2 Analyses

For each task, a population-level dataset was created using all available participants, which we refer to as the *gold standard*. Studies were generated by drawing bootstrap samples (with replacement) from the gold standard. These studies are referred to as *generated studies* or *bootstrap samples*. For the HCP dataset, study sample sizes were 20, 40, 60, 80, or 100 participants. For UKB, study sample sizes were 20, 40, 60, 80, 100, 1000, and 10000. At each sample size, 100 bootstrap samples were generated.

#### 2.2.1 Region of Interest

Voxels or vertices were labeled and grouped according to either the Schaefer parcellation (after projection to standard space) or the Harvard-Oxford subcortical atlas (Schaefer et al., 2018). Analyses were performed using the 400-level parcellation (additional levels are presented in the Supplementary Materials). Regions were further assigned to one of the Yeo7 Networks or, if they were within the subcortex, labeled as such (Thomas Yeo et al., 2011).

Region-of-interest analyses were based on both binary activation maps and regional effect size maps (Cohen’s *d*). To calculate activation maps, each subject’s average activity was computed per region, and regional group activation was tested using a *t*-test across participants at *α* = 0.05 after family-wise error correction using Holm correction (Holm, 1979).

To facilitate comparisons across tasks, validity analyses focused on the ten regions that exhibited the largest effect size in the gold standard for each task. To compare the generated studies to the gold standard, we calculated either the proportion of studies at each sample size that exhibited significant activation in each of these ten regions (binary activation maps) or the rank correlation between the study’s regional effect sizes and the gold standard’s regional effect sizes (continuous effect size maps).

To assess the reliability of the generated studies, we calculated intraclass correlation coefficients (ICCs) across bootstrap samples. For binary activation, the ICCs were estimated using a Monte Carlo method (1000 samples) implemented in the aod package (Lesnoff et al., 2012), which is based on a one-way random effects model (Goldstein et al., 2002), and confidence intervals were estimated via percentage bootstrap sampling (100 samples). For analyses of regional effect sizes, the ICCs were calculated using the irr package (Gamer et al., 2019), using a single unit, two-way random effects measure of consistency: ICC(C,1). In both cases, ICCs were computed over the complete set of regions. Restricting to the top ten regions yielded degenerate estimates in the binary case due to near-zero variability across bootstraps because all studies were often significant.

#### 2.2.2 Peak Localization

Peaks for the gold standard were extracted based on the raw group-level *t*-statistic map derived from all participants. In the volumetric analyses, a peak was defined as any voxel larger than all of its 26 connected neighbors, calculated using cluster from FSL (S. M. Smith et al., 2004). In the surface-based analyses, a peak was defined as a vertex exceeding all other vertices (or voxels, in the case of subcortex) within a distance of 1 mm.

Peaks in sampled studies were extracted from unsmoothed, threshold-free cluster-enhanced *z*-maps (S. Smith & Nichols, 2009) after family-wise error-corrected thresholding at *p <* 0.05, calculated using permutation tests (Winkler et al., 2014). This thresholding removed voxels with negative activation, so peak location displays only include positive effects. Given the computational demands of permutation testing, peak localization analyses were limited to studies with sample sizes of 40 and below. Exploratory analyses with volumetric data using probabilistic threshold-free cluster enhanced *z*-maps (Spisák et al., 2019), which does not require permutation testing but is not yet available for surface meshes, indicated that results stabilize by 40 participants (Figure S2). Unthresholded results are presented in the Supplementary Material.

Analogously to the region of interest analyses, we sought to facilitate comparisons across tasks by considering only the subset of peaks that are most relevant for each contrast. Specifically, we considered the ten peaks that had the largest (positive) activation in the gold standard, taking at most one peak per parcel or region (the largest). These were compared to the sampled studies by calculating the proportion of such studies that contained any local peak within different radii (2, 4, 8, 10, 20 mm) of a sphere centered at each gold standard peak.

To compare between studies, we determined the peaks from the *z*-maps of individual studies or the *z*-maps from individual participants that were closest to the ten gold standard peaks. Comparisons were made by assessing the distributions of peak distances. In volumetric analyses (including those involving the subcortex for SURFACE and MSMAll analyses), distances were calculated as the Euclidean distance between voxel coordinates in the reference space. In surface-based analyses, distances were computed in Connectome Workbench using the non-naïve method (Marcus et al., 2011).

#### 2.2.3 Topography

Voxel-wise effect sizes were computed using Cohen’s *d*. To construct the gold standard map, for voxel *v*, effect sizes were calculated using the across-subject mean *µ*_*v*_ and standard deviation *σ*_*v*_ of each contrast of parameter estimate (i.e., 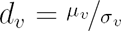). Voxels were binned into categories using the guidelines provided by Cohen (1988): with “negligible” indicating |*d*_*v*_| *<* 0.2, “small” indicating 0.2 ≤ |*d*_*v*_| *<* 0.5, “medium” indicating 0.5 ≤ |*d*_*v*_| *<* 0.8, and “large” indicating 0.8 ≤ |*d*_*v*_|. The effect sizes in sampled studies were calculated using Hedges’ small-sample bias correction (Bossier et al., 2019; Hedges, 1981).

To compare the gold standard with the generated studies, rank correlations were calculated between the maps of the gold standard and the sampled studies, within a gray matter mask. To assess the reliability of sampled studies, we calculated ICC(C,1) intraclass correlations across bootstrap samples.

#### 2.2.4 Multivariate Models

Models predicting participant traits were trained using task-based functional connectomes. For both datasets, the data were subjected to basic cleaning: linear detrending, band-pass filtering (0.01–0.1 Hz), voxel-wise standardization, and nuisance regression with 24 motion parameters (Friston et al., 1996; Satterthwaite et al., 2013). To build connectivity matrices, the timeseries for each task (after concatenation of runs, in the case of HCP-YA) were parcellated using the Schaefer100 atlas (Schaefer et al., 2018). Connectivity was estimated using Ledoit–Wolf shrinkage of the covariance matrix (Ledoit & Wolf, 2004), and the resulting pairwise correlations were Fisher transformed with the inverse hyperbolic tangent. Features consisted of the 4950 lower-triangular elements of the connectivity matrix. Outcomes consisted of instruments such as task performance and fluid intelligence. For a complete list, see (Table S2).

Considering the small sample sizes and large numbers of features, models relied on feature selection and regularization implemented in scikit-learn (Pedregosa et al., 2011). First, features with variance less than 0.01 were removed. Then, features were independently standardized using robust normalization (i.e., removing the median and scaling by the interquartile range). Predictions were made using ridge regression with the regularization parameter selected from 20 log-spaced values (0.1 to 10,000, inclusive) using the efficient leave-one-out cross-validation procedure described by Rifkin and Lippert (2007) and implemented in scikit-learn. All preprocessing and screening were fit within training folds and applied to held-out data to prevent information leakage.

To facilitate comparisons across study sizes (and with the gold standard), we held out a fixed test set consisting of 20 % of the full dataset from all training and validation. The same test set was used for all sampled studies of a given task, and did not include any participants from the gold standard. That is, while the model hyperparameter (regularization parameter) was determined using leave-one-participant-out cross-validation, the final model performance was measured on a group of participants whose data did not contribute to model training or hyperparameter tuning.

We considered three kinds of modeling endpoints: model performance, prediction significance, and model coefficient weighting. Model performance was measured as the rank correlation between trained model predictions and the true values in the test set (see also Supplementary Materials for assessment by *R*^2^, Figure S10). The gold standard was defined as the model’s performance on the test sample when the training sample comprised all participants except the ones being evaluated. This gold standard was compared to the performance of the sampled studies at each sample size. To compare performance across studies, the predictions on the held-out test set were used to calculate an intra-class correlation based on a two-way random-effects model. Given that we were interested in the reliability of a single study, we used the single-measure version, resulting in a measure known as ICC(C,1) (McGraw & Wong, 1996). Intraclass correlations were calculated using the R package irr (Gamer et al., 2019; R Core Team, 2023).

The validity of prediction significance was measured analogously, except that studies were summarized according to whether the rank correlation on the test set was statistically significant, using permutation tests on the rank correlations, each with 1000 permutations. The intraclass correlation was not calculated for rates of significance because it is not meaningful for binary outcomes. There is only a single outcome of significance for each simulation (on the held-out set), and thus, there is insufficient data to measure within-simulation variability. Instead, we report on the variability of the generated datasets (variability of a Beta distribution estimated using analytic Bayesian methods and Jeffrey’s priors). Validity of model coefficients was assessed with the distribution of product-moment correlations between the coefficients of the gold-standard model and the study-trained models, and reliability was evaluated by comparing study-trained model coefficients to each other using ICC(C,1).

## 3 Results

### 3.1 Region of Interest

#### 3.1.1 Gold Standard

We evaluated statistical power to detect regional activation as a function of sample size. For each task, we designated a set of regions as “primary targets” if their average effect size in the gold standard was among the ten highest across the brain (Table S4). This designation enabled us to focus on a core set of regions for each task, representing regions that were likely to be studied in conjunction with the given task (for region definition, see Methods). For example, in the motor task, this approach picks out voxels within the primary motor cortex. In assessing validity and reliability, we considered both a binary version of activity (thresholded by statistical significance) and the raw effect size.

#### 3.1.2 Study Validity

Studies based on HCP-YA using the emotion, language, and motor tasks were nearly guaranteed to detect effects in most of the ten targeted regions, even with only 20 participants (Figure 2a). In contrast, for the UKB emotion task, a subset of the top regions did not reach maximal significance rates with fewer than 100 participants (compare these results with those for VOL in HCP-YA). In the HCP-YA social and working memory tasks, similarly high power would require around 40 to 60 participants, with the volumetric analyses requiring larger sample sizes. Finally, the gambling and relational tasks would need between 60 and 80 participants. These patterns were largely consistent across different levels of parcellation granularity (Figure S3).

**Figure 1:**
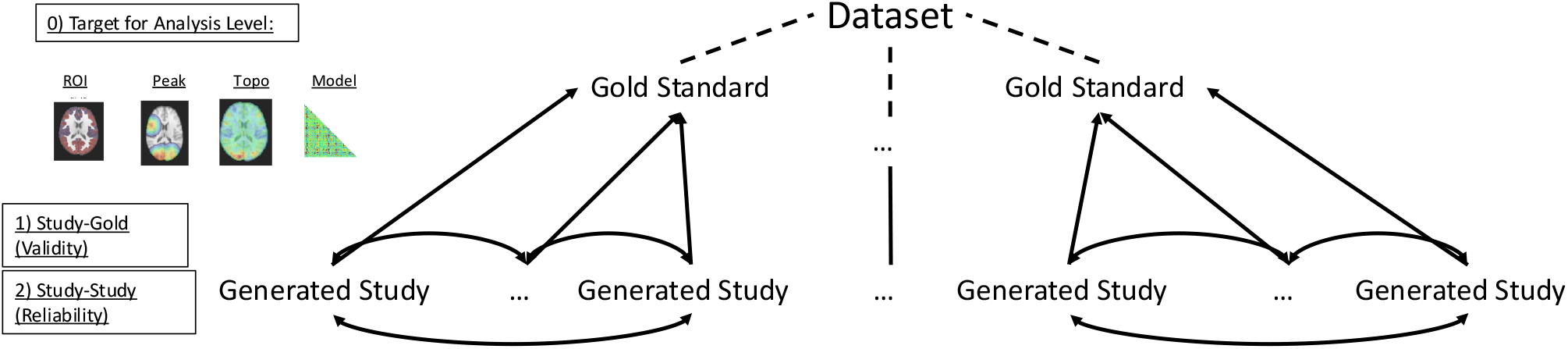
Outline of Methods. Using two large datasets (HCP-YA and UKB), we constructed gold standards at each of four different analysis levels: region of interest activation (subsection 3.1), peak localization (subsection 3.2), topography (subsection 3.3), and predictive model performance (subsection 3.4) by applying identical pipelines to the full cohorts. Smaller studies were generated by bootstrapping participants at target sample sizes, repeating the same analyses, and comparing the results to the gold standards and across studies.

**Figure 2:**
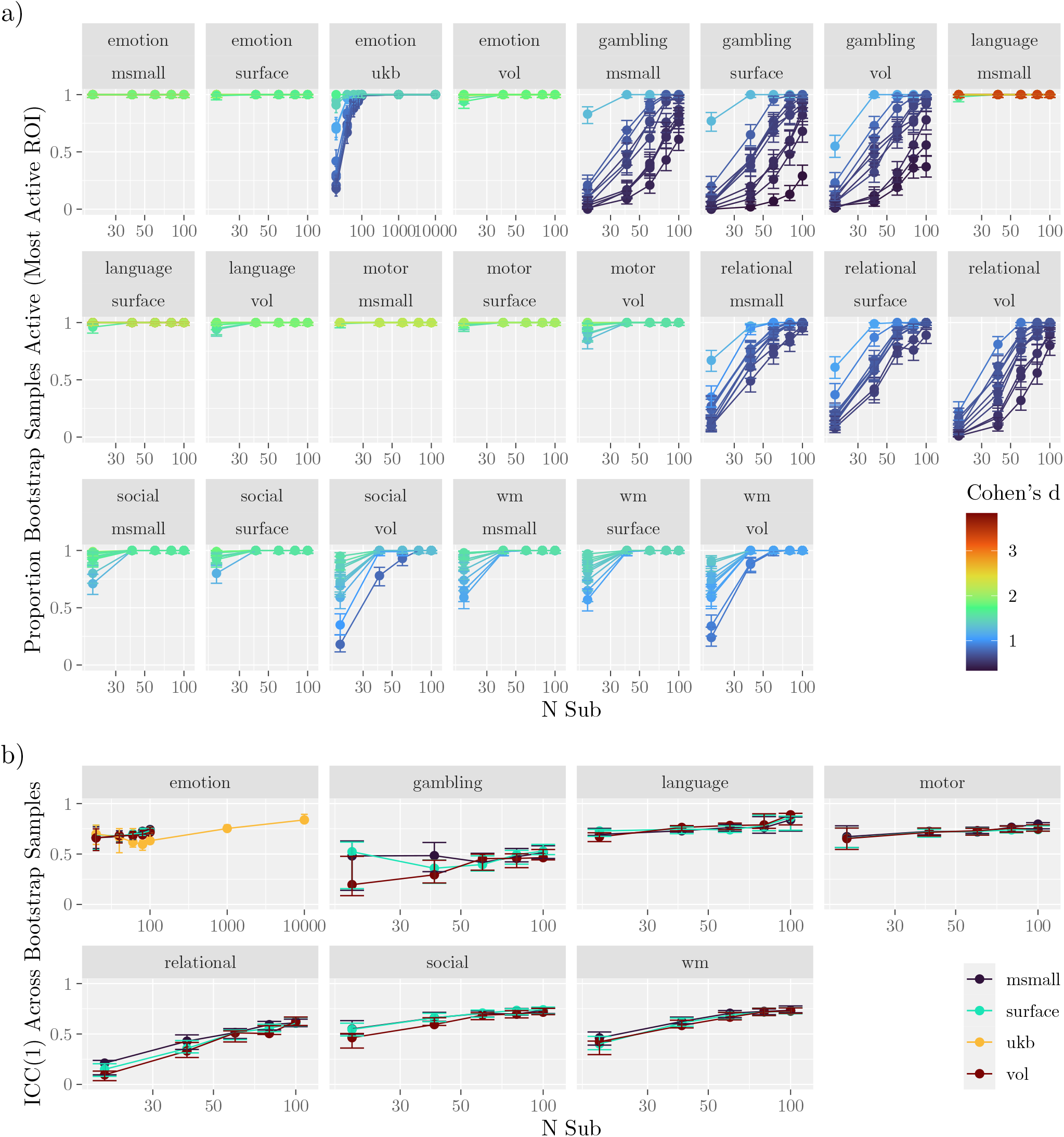
Activation Within Regions of Interest. **a)** Validity: Proportion of bootstrap samples that show activation in each of the ten ROIs with the largest gold-standard effect sizes. Results are plotted across a range of sample sizes. Each line corresponds to one of the ten ROIs. Error bars span the 95 % highest-density interval. **b)** Reliability: Consistency across bootstrap samples measured using the intraclass correlation coefficient (ICC) as a function of sample size. Note that the ICC was calculated using all regions.

Validity of the unthresholded maps followed a similar pattern (Figure 3a). The language task exhibited correlations with the gold standard in excess of 0.9, even with only 20 participants. The motor task exhibited similarly high correlations on average (minimum of 0.765). The other tasks also exhibited high average correlations, albeit with wider confidence intervals, highlighting larger variation across the generated studies. For example, the interval for the working memory task included 0. The confidence intervals for each modality (VOL, SURFACE, MSMAll) overlapped.

**Figure 3:**
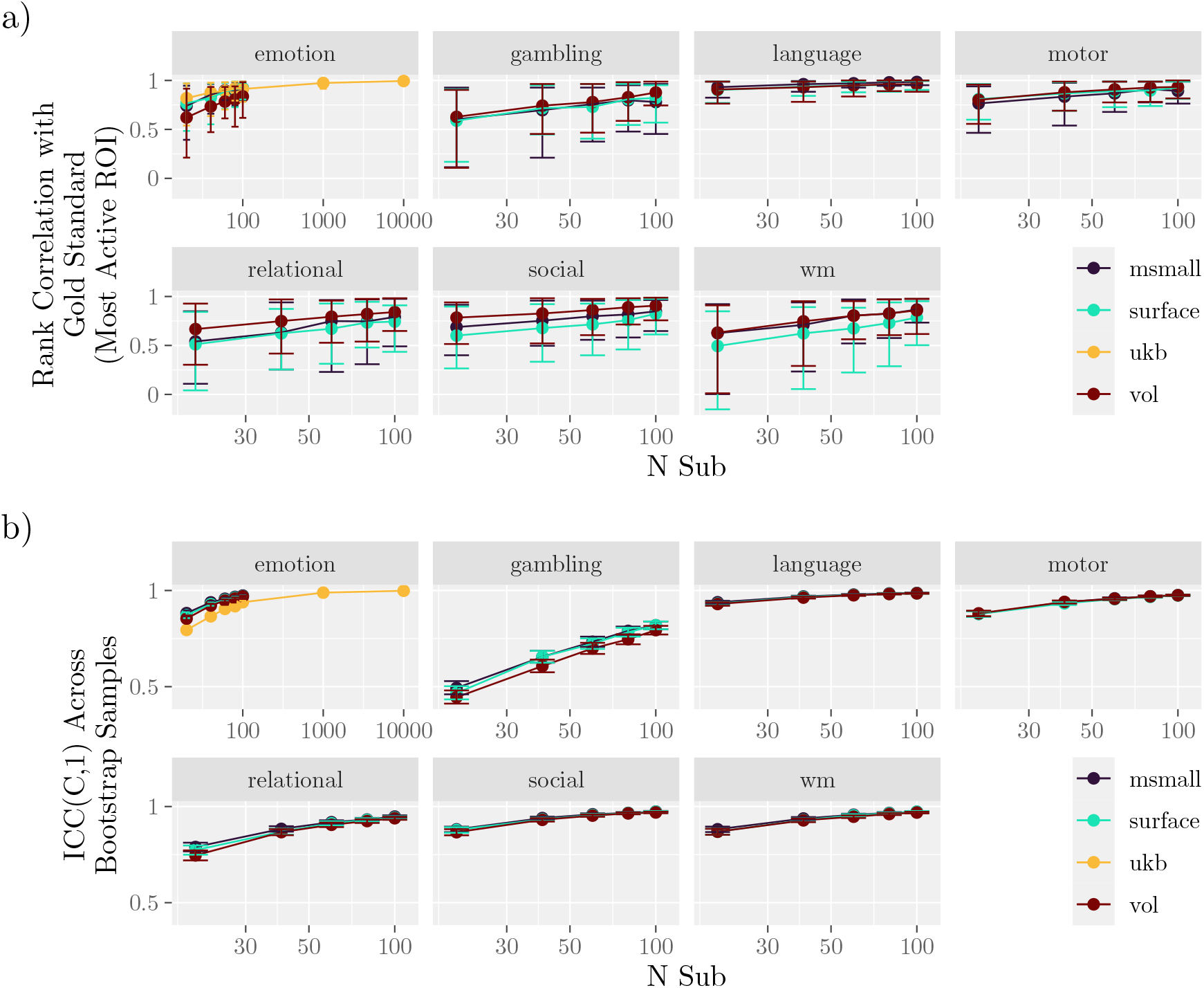
Effect Sizes Within Regions of Interest. **a)** Validity: The rank correlation between the bootstrap samples and the gold standard regional effect sizes. Results are shown for three data types from the HCP-YA cohort (VOL, SURFACE, MSMALL) and volumetric data from the UKB. Error bars span the 2.5 % to 97.5 % quantiles across samples. **b)** Reliability: Consistency across bootstrap samples measured using the intraclass correlation coefficient (ICC) as a function of sample size. Error bars span 95 % confidence intervals. Note that the intraclass correlation was calculated using all regions.

#### 3.1.3 Study Reliability

To assess the reliability of regional significance, we first calculated the intraclass correlation of the activation vectors for each of the generated studies (Figure 2b). Tasks with the most reliably activated primary target regions showed the highest coefficient distributions. Language, motor, and emotion tasks consistently exhibited an intraclass correlation that was “moderate” to “good” (Koo & Li, 2016). The relational, social, and working memory tasks achieved similar levels after samples included 40 to 60 participants. Even with 100 participants, the intraclass correlation for the gambling task was still only around 0.5.

Across sample sizes, the consistency of regional effect sizes was nearly perfect for the language, motor, relational, social, and working memory tasks (Figure 3b). Consistency in the gambling task was the lowest. Differences between modalities were minimal for most tasks (e.g., SURFACE and MSMAll were nearly indistinguishable).

### 3.2 Peak Localization

#### 3.2.1 Gold Standard

Next, we examined peak localization. Because large sample sizes produced activation clusters spanning much of the gray matter, we focused on local rather than global cluster peaks. As before, we assume that each task induces activation in multiple distinct regions, with the number of regions varying between tasks (see Table S3 for a list of regions). To facilitate comparisons between tasks, for each contrast, we selected the ten highest local maxima (not necessarily the same regions exhibiting the largest effect sizes).

#### 3.2.2 Study Validity

To quantify validity independently of the atlas used, we measured the distance between the highest gold-standard peak and the nearest peak within each study. Specifically, we calculated the proportion of sampled studies that contained a significant peak within various radii, plotting these proportions by task and sample size (Figure 4). Increasing the sample size improved peak localizability (compare results across columns of Figure 4), although for many regions, validity was not substantially higher for studies with more than 40 participants (Figure S2). In all tasks except gambling and relational, nearly all generated studies produced peaks within 10 mm of the gold standard peaks — even with only 20 participants. For the gambling and relational tasks, studies with 20 participants often failed to provide a peak within that radius. Even so, with 40-60 participants in these two tasks, over 95 % of studies produced peaks that were within 10 mm of one of the top ten peaks (Figure S2), equivalent to five voxels in the HCP-YA dataset.

**Figure 4:**
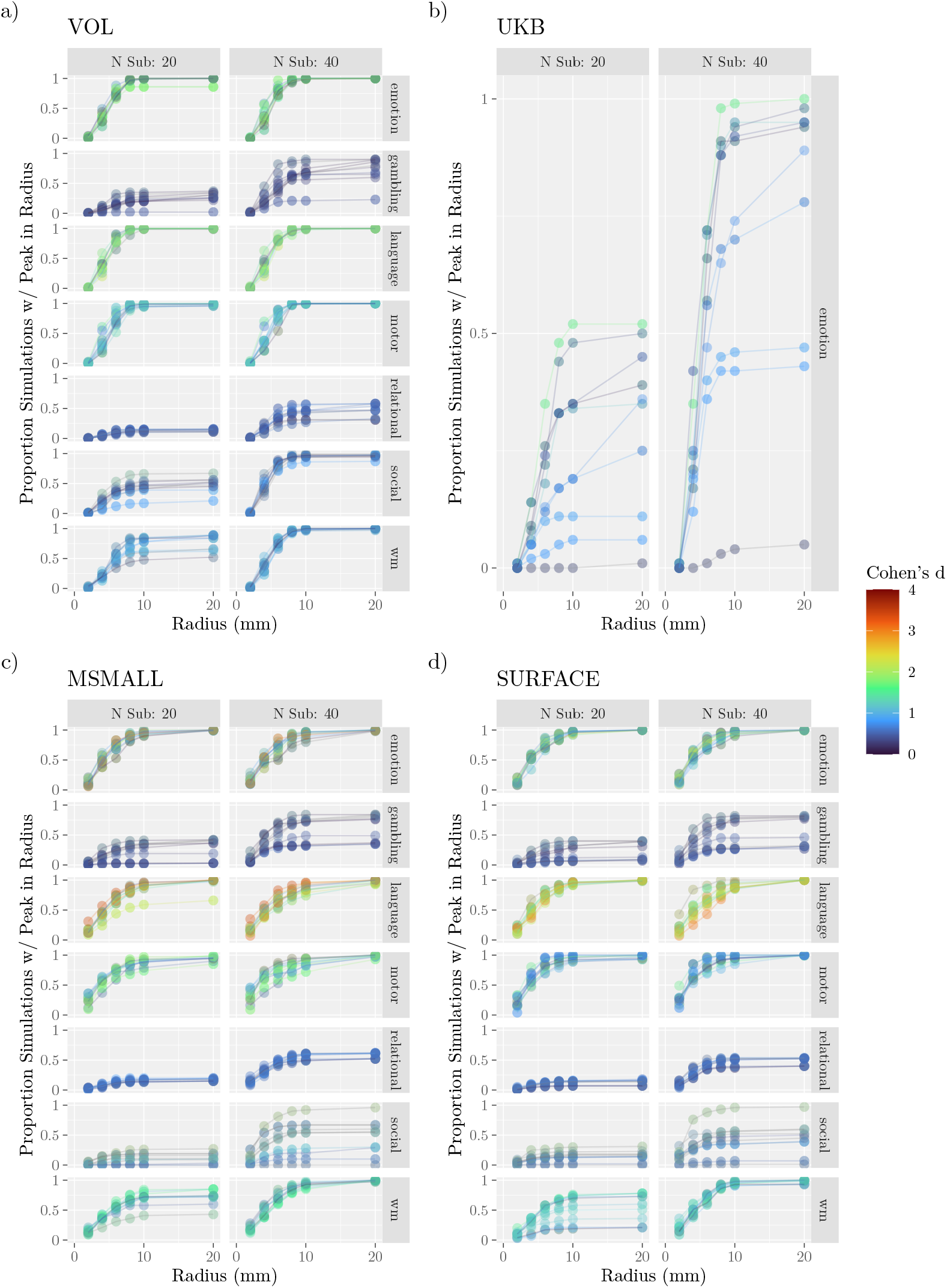
Localization of Peak Activation. Validity: Proportion of studies containing peaks within a given radius. Rows indicate tasks from the HCP-YA dataset, and columns are the sample sizes. Each panel depicts the proportion of studies that contained peaks that were within a given radius of one of the 10 largest peaks for that task, with peaks colored by the effect size of that voxel in the gold standard. The figure considers only supra-threshold peaks (compare with Figure S4). Results shown for (a) HCP-YA volumetric data (VOL), (b) UK Biobank volumetric data, (c) HCP-YA MSMAll data (MSMALL), and (d) HCP-YA surface data (SURFACE).

When considering all peaks per contrast (i.e., those with non-negligible effect sizes in voxels that survived family-wise error correction), localization depended strongly on effect size (peaks with larger effects in the gold standard were better localized with fewer participants) (Figure S7, see also Roels et al., 2015). For example, in generated studies of 20 participants, peaks in the gold standard with small effect sizes (|0.1 *−* 0.3|) were separated from the study peaks by an average of 40 mm, while the same average for peaks whose voxels had a large effect *>* |0.5| was only 11 mm.

There were apparent differences in localizability when categorizing peaks according to connectivity network (Thomas Yeo et al., 2011), such that peaks within “lower-level” networks, such as somatosensory or visual networks, were localized more easily than those within “higher-order” networks like the default or limbic networks (Figure S6). However, there was also a close relationship between the presence of a peak within a network and the height of the peak, so the effect of the network was not necessarily distinct from the impact of peak height. For example, consider that, with only 20 participants, peaks within the somatomotor network were an average of 8.5 mm from the gold standard in the motor task (average effect size: 0.41), but for the emotion task that distance jumped to 42.5 mm in the social task (average effect size: 0.16).

Note that all of these counts are conditional on the presence of a peak and that not every generated study produced a supra-threshold peak. For the number of generated studies without peaks, see Table S1.

Finally, while Figure 4 suggests substantial differences between UKB and HCP-YA datasets for an analogous task (emotion), the magnitude of this difference depends strongly on thresholding; when considering unthresholded maps, a much higher proportion of studies with UKB participants had peaks that were within 10 mm of the gold standard peaks (Figure S4). A similar strong dependence on thresholding was also observed for the worst-performing tasks in the HCP-YA dataset (gambling, relational).

#### 3.2.3 Study Reliability

To compare studies, the local peaks associated with the ten highest peaks were grouped, and the pairwise distances between them were calculated (Figure 5). These distances highlight the expected variability across studies with small sample sizes. For example, with 20 participants, 10 % of gambling studies had peaks that were over 19.7 mm apart in the VOL analyses. In contrast, for the same sample size and modality, the 90th percentile for the motor task was only 4.69 mm. Note that these differences are highly dependent on the significance of the peaks; without thresholding, the 90th percentiles for gambling and motor were 5 mm and 4.47 mm.

**Figure 5:**
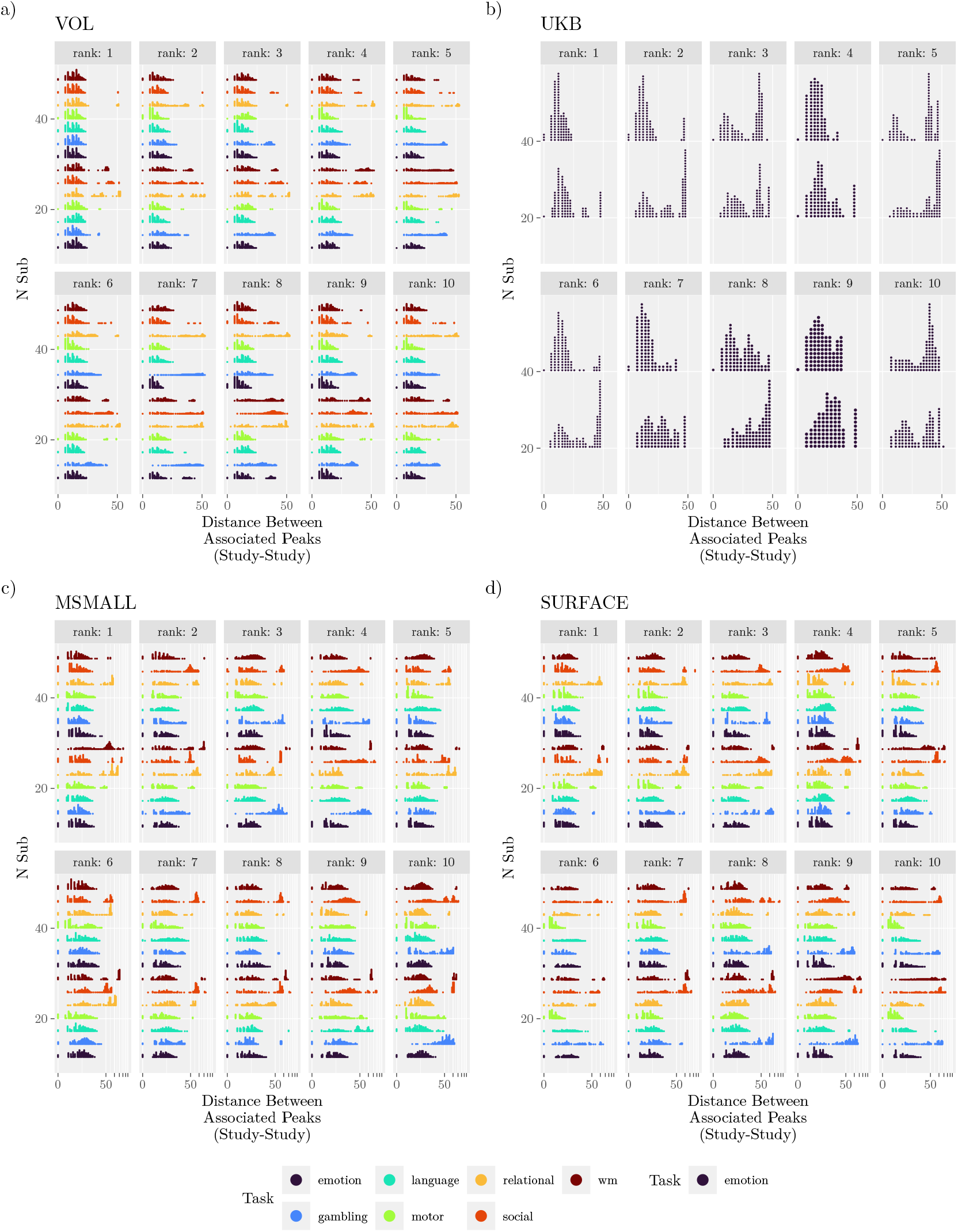
Localization of Peak Activation. Reliability: Average distance between peaks in studies that were associated with common peaks in the gold standard. Within distributions, points represent percentiles of distances. The figure considers only supra-threshold peaks (compare with Figure S5). Results shown for (a) HCP-YA volumetric data (VOL), (b) UK Biobank volumetric data, (c) HCP-YA MSMAll data (MSMALL), and (d) HCP-YA surface data (SURFACE).

### 3.3 Topography

#### 3.3.1 Gold Standard

Peaks capture only one aspect of activation, so we next examined voxel- or vertex-wise effect sizes. The contrasts for the gambling and relational tasks had distributions with the smallest averages, resulting in the largest proportion of voxels with negligible effects and the smallest proportion of voxels with medium and large effects (Figure S8). For the gambling task, fewer than 1 % of voxels had an effect size that was medium or large. In the language, motor, social, and working memory tasks, small effects were present in around 30 to 40 % of voxels, and medium effects were present in 10 to 25 %. In most tasks, large effects were present in fewer than 5 % of voxels. But in the language task, a large effect was present in almost 20 % of voxels. Trends were similar in vertex-based analyses (Figure S8).

#### 3.3.2 Study Validity

For all tasks, 99 % of rank correlations between the effect size maps of the studies and the gold standard map were above 0.5 (Figure 6a). Consistent with the gambling task eliciting smaller effects, the correlations for this task were generally lower. In contrast, the language task, which tended to elicit the strongest activation, showed correlations that were typically above 0.75 at all sample sizes.

**Figure 6:**
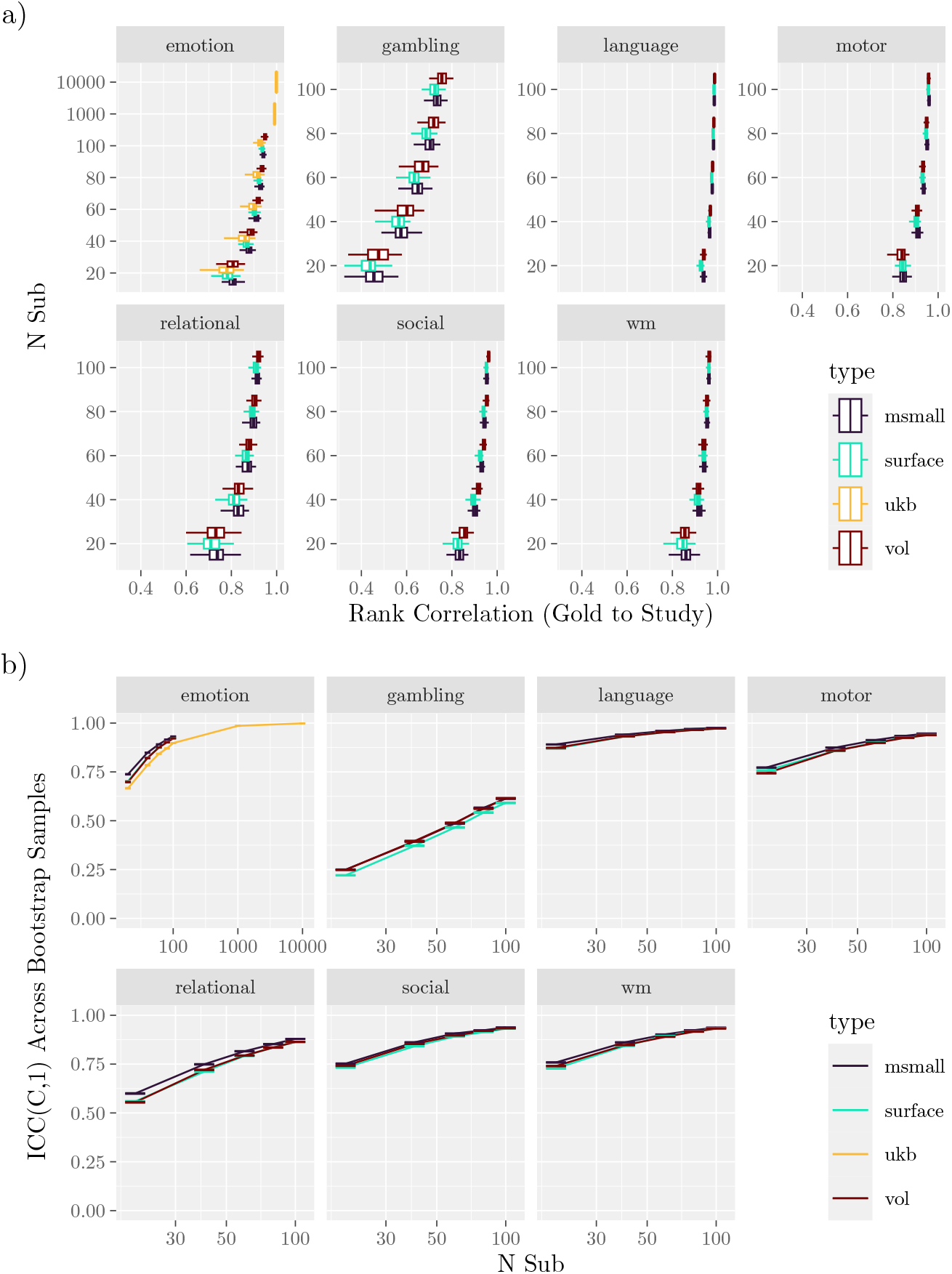
Topographic Maps. **a)** Validity: The rank correlation between the gold standard and bootstrap samples generated under a range of sample sizes. **b)** Reliability: Rank correlations were calculated between the gold standard map and the maps from the bootstrap samples. Comparisons between bootstrap samples were performed using the intraclass correlation coefficient (ICC) for various sample sizes. The correlation was calculated with each spatial point (vertex or voxel) as a class. Error bars span 95 % confidence intervals.

As with the reliability of activation and peak localization, there was variation in the recovery of the gold standard across networks, and that variation was consistent with an important role for the effect sizes within the networks (Figure S9). Across all tasks, voxels within the limbic network were among those that exhibited the lowest correlations. In most tasks, voxels within the subcortical regions also exhibited low correlations. Correlations for voxels within the somatomotor network were neither the highest nor the lowest for all tasks except the motor task, where they were the highest.

#### 3.3.3 Study Reliability

Regarding reliability, the gambling task exhibited the lowest correlations, ranging from approximately 0.25 with 20 participants to 0.55 with 100 participants. In contrast, the language task exhibited the highest correlations, ranging from around 0.9 with 20 participants to.97 with 100 participants. As in comparisons of study validity, there were no substantial differences in the reliability of effect sizes across data types.

### 3.4 Multivariate Models

#### 3.4.1 Gold Standard

Models were trained to predict characteristics related to cognition in the HCP and UKB datasets. Predictions were based on features derived from connectivity matrices and a ridge regression model.

With the gold standard, there was substantial variability in model performance across instruments and tasks (Figure 7a, Figure S10). Tasks such as working memory and language enabled the model to achieve correlations on the held-out dataset exceeding 0.25 for several instruments. In contrast, the motor task achieved such high correlations for only a few measures. When training with the full dataset, a subset of HCP instruments exhibited negative correlations on all tasks (different instruments for each task). For a complete list of performance on each task, see the Table S2.

**Figure 7:**
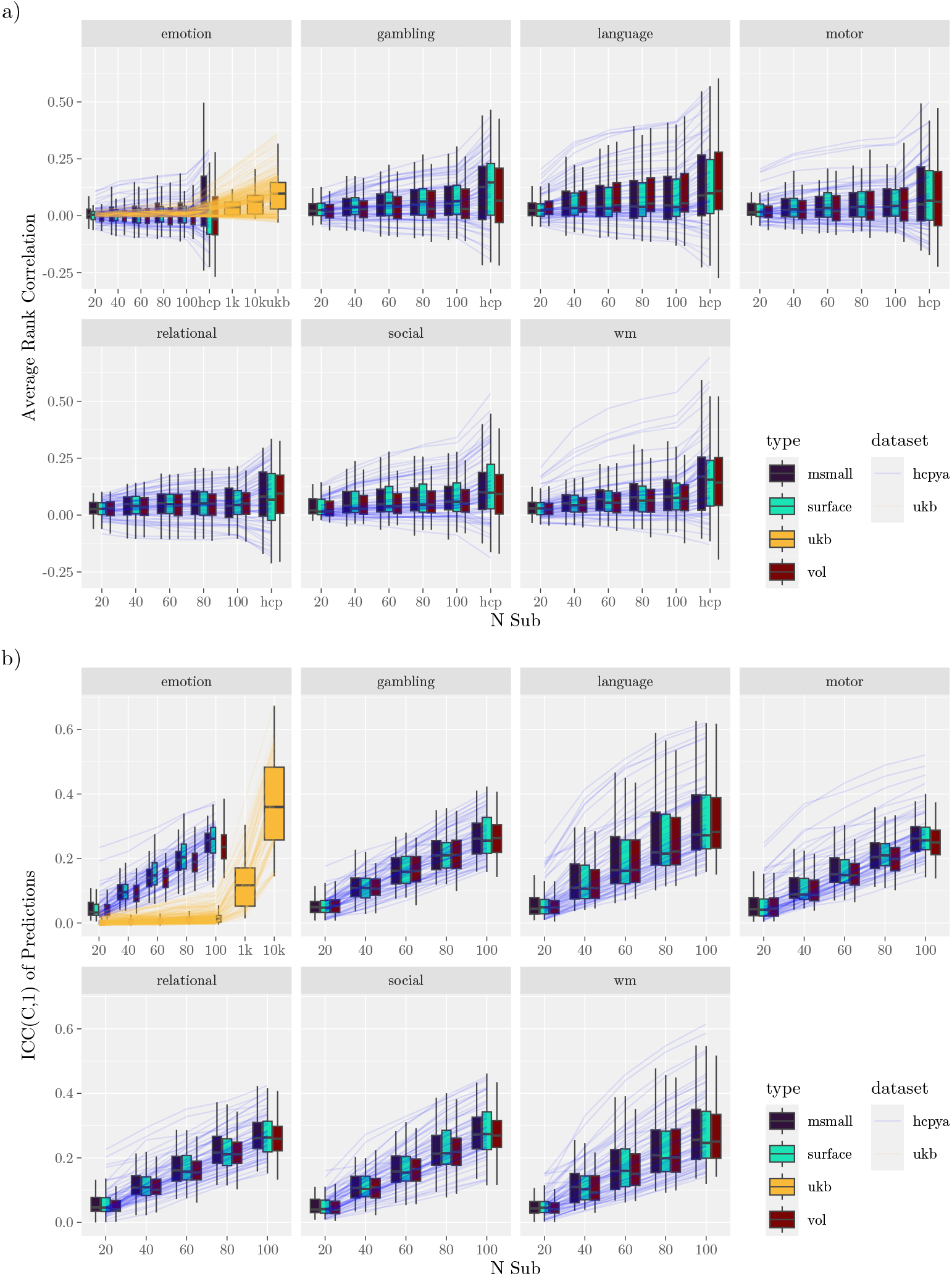
Multivariate Pattern Models Prediction Scores. **a)** Gold standard and comparison between the gold standard and the generated datasets. The lines trace the rank correlation between model predictions and the true values in held-out samples (average correlation). For performance as measured with the coefficient of determination, see Figure S10. **b)** Reliability of model predictions across samples. In both subfigures, the lines correspond to different measures (averaged across type within each dataset).

In a supplementary analysis, model performance is measured with a different feature set: not the connectome but instead the effect sizes from the ten regions that were most active (Figure S11). The trends in performance were similar, but overall performance was lower (e.g., no measure was predicted by the gold standard with a correlation above 0.2).

Regarding statistical significance, the HCP emotion task supported significant predictions in the smallest subset of features (14 % compare Figure 8 and Figure 9), and the working memory and language tasks supported the largest subset (38 %). The UKB emotion task supported significant predictions in 75 % of instruments (instruments differed between the HCP and UKB).

**Figure 8:**
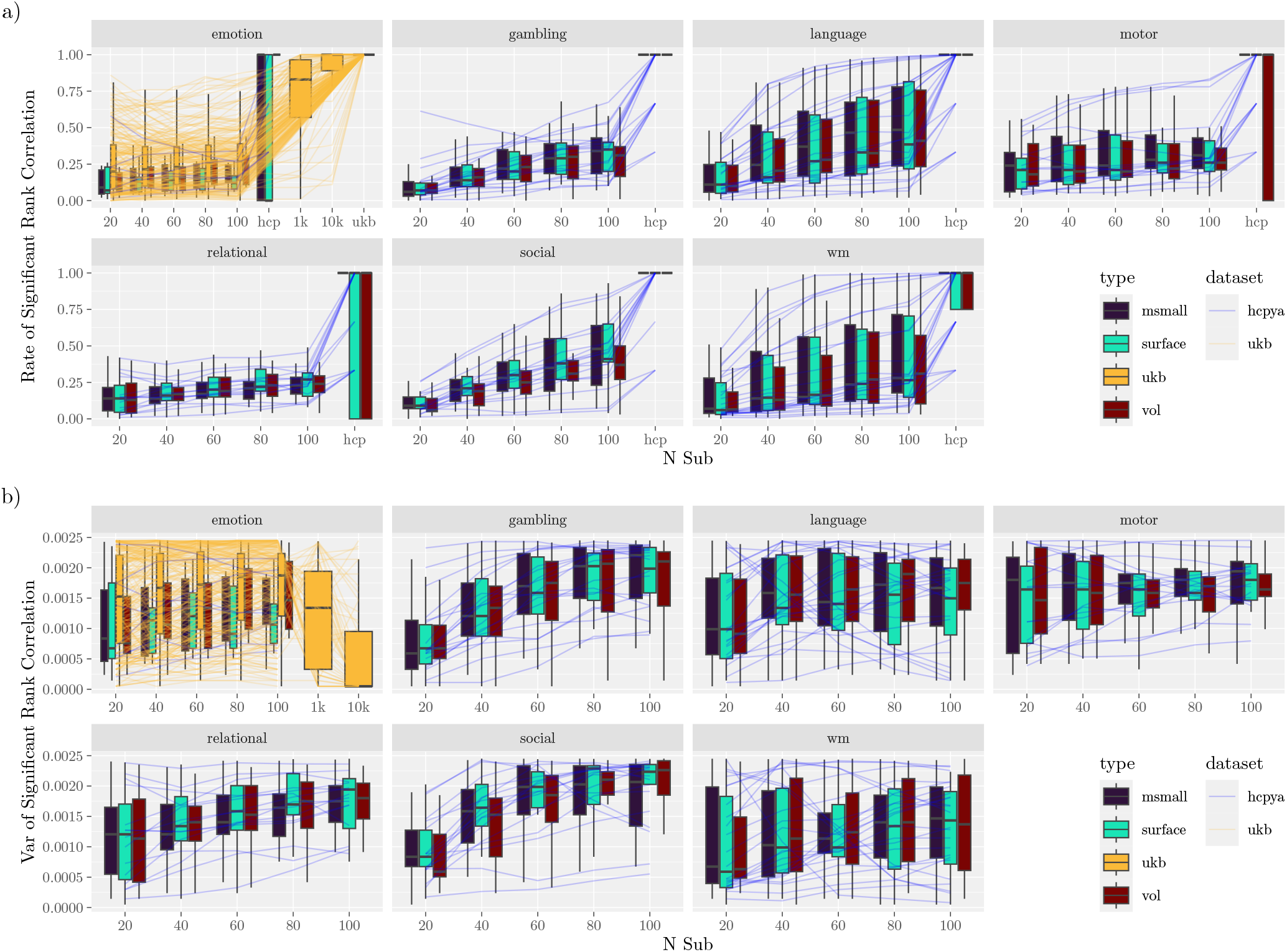
Multivariate Pattern Models Prediction Significance. **a)** Gold standard and comparison between the gold standard and the generated datasets. Lines correspond to different measures (averaged across type within the dataset), limited to measures that were significant across all types in the gold standards. Higher values correspond to lower false negative rates (cf., Figure 9).

**Figure 9:**
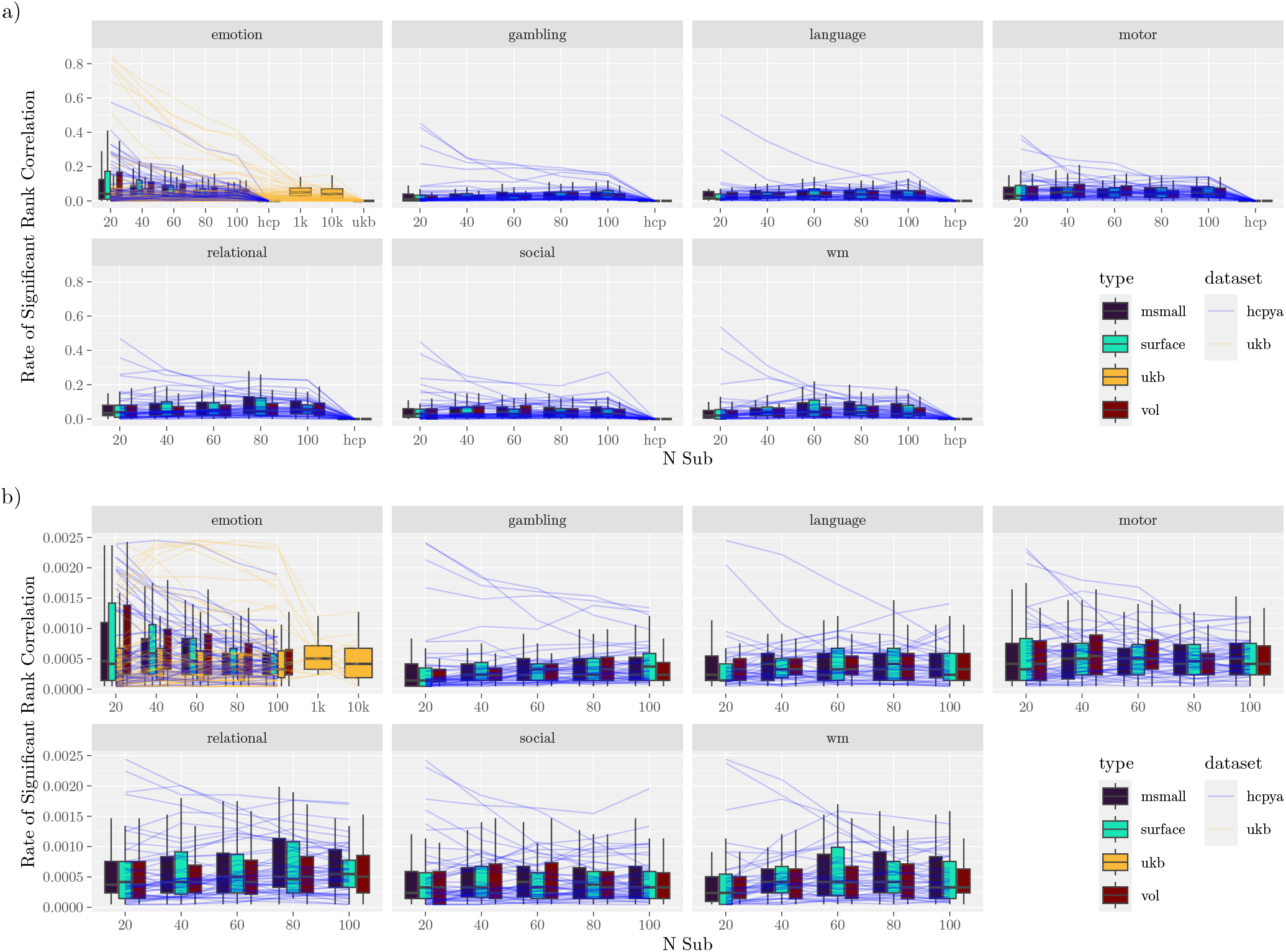
Multivariate Pattern Models Prediction Significance. **a)** Gold standard and comparison between the gold standard and the generated datasets. Lines correspond to different measures (averaged across type within the dataset), limited to measures that were not significant in all types in the gold standard. Lower values correspond to lower false positive rates (cf. Figure 8).

#### 3.4.2 Study Validity

Study validity was first assessed by comparing the levels of predictive performance achieved in the generated studies with those achieved using the gold standard. All tasks exhibited performance that was numerically below the gold standard level, even with the highest sample sizes considered (100 for the HCP and 10,000 for the UKB, Figure 7a, Figure S10). Model performance increased steadily as the sample size increased.

Next, we examined the rate at which individual studies provided significant model performance (Figure 8 and Figure 9), which can be considered an estimate of statistical power (Figure 8) or false positive rates (Figure 9). For most tasks and instruments, power was well below the standard 80 %, even with 100 participants. The only exception to this was the language and working memory tasks, which allowed greater than 80 % power for 6/24 and 5/24 instruments, as well as the UKB emotion task, which provided similarly high power for 3/220 instruments.

Finally, we assessed how well individual studies estimated model features (Figure 10). In the HCP dataset, correlations ranged between 0.2 and 0.5, steadily increasing with sample size. In comparison to the HCP, feature recovery was worse at lower sample sizes, remaining below 0.2 with fewer than 1,000 participants, and only matching the HCP at the largest sample sizes considered (10,000 participants). There were only minimal differences in feature recovery across tasks and data types.

**Figure 10:**
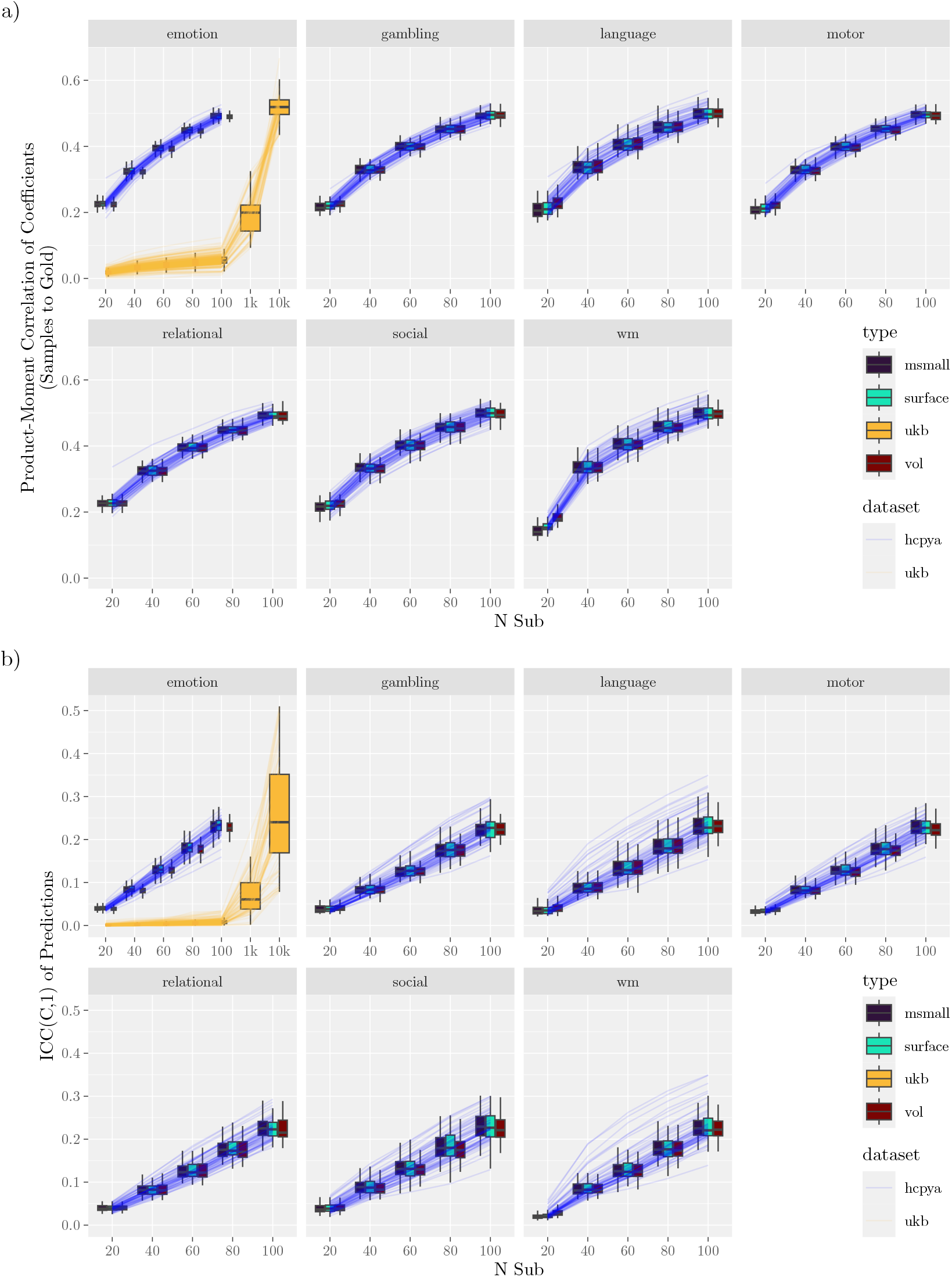
Coefficients for Multivariate Pattern Models Prediction Significance. **a)** Gold standard and comparison between the gold standard and the generated datasets. Lines trace the average correlation between features in the model estimated with the gold standard dataset and features in the study datasets. **b)** Reliability of model predictions across samples. In both subfigures, the lines correspond to different measures (averaged across type within each dataset).

#### 3.4.3 Study Reliability

To assess study reliability, we first calculated the intraclass correlation for participant-level predictions (Figure 7b). This enables asking about the stability of predictions for a given subject across training datasets: by how much do predictions vary according to the training set? Across all tested sample sizes and tasks, reliability was poor (Koo & Li, 2016), with intraclass correlations below 0.5 (Figure 7a). At a sample size of 100, only the language, working memory, and motor tasks supported predictions with ICCs above 0.5 (rates of instruments: 7/63, 4/63, 1/63). The UKB data required sample sizes of at least 10,000 to achieve intraclass correlations above 0.5 (17/74).

Reliability was equally poor for feature reliability (Figure 10b). With 1000 participants, no measure had an ICC greater than 0.5. With 10000, 72 % of measures were above 0.5, but none indicated “good” reliability (¿0.75 Koo & Li, 2016).

## 4 Discussion

In this report, we leveraged the Human Connectome Project and UK Biobank datasets to survey several aspects of validity and reliability in group-level task-based fMRI studies conducted with sample sizes typical in the neuroimaging literature — often less than 100. Using these rich datasets, we first constructed gold standards based on the complete datasets. We then explored how well features of those standards could be recovered by smaller studies (validity), and the extent to which smaller studies were consistent with each other (reliability). Our goal was to help researchers calibrate their expectations about the informativeness of individual studies by considering the influence of sample size, effect size, and the choice of analysis.

First, we considered the classical mass-univariate approach for detecting activation in task-related regions. Predictably, regions with large effect sizes could be detected with relatively small sample sizes — around 40 participants (Figure 2a). For context, achieving 80 % power with a one-sample, two-sided *t*-test with a true effect size of 0.8 requires at least 15 observations, which increases to 34 observations with an effect size of 0.5, and with an effect size of 0.2, 199 observations are required. However, large effect sizes were uncommon and nearly absent from some tasks (Figure 6a, Figure S8). This means that reports of novel effects – even large ones – based on only 40 participants should be interpreted with caution (see also Marek et al., 2019; Reddan et al., 2017). While observing a significant effect with 40 participants can justify continuing a line of research, the scarcity of large effects underscores the need for replication before a finding is considered established.

When examining peak activation, we observed that studies with 40 to 60 participants were generally able to localize a peak to within 10 mm of the gold standard. This result held across both volumetric and surface-based analyses (Figure 4). To put this in perspective, the estimated number of distinct regions within the cortex is around 300 - 400 (Van Essen et al., 2012). At that scale, spherical regions would have a radius of around 8.3 mm, suggesting that with 40 participants, a study’s local peaks are likely to fall within one to two regions of the gold standard peak. While this resolution may be inadequate for fine-grained anatomical questions (e.g., segmenting subcortical microstructures), recall that a 10 mm distance translates to only a few voxels in these datasets.

Variability in peak localization can be interpreted in several ways. The true peak activation location (voxel or vertex) may be the same across all individuals. In this case, variability across bootstrapped studies would reflect variance in the ability to detect that peak within samples. There is some evidence for such variability; across repeated scans of the same individual, estimated cluster center-of-mass may vary by around 1.8 mm to 3.9 mm (Marshall et al., 2004; Rath et al., 2016; Rombouts et al., 1998; Wurnig et al., 2013), although that variability increased when scans are collected at different sites (Rath et al., 2016). In a patient population, peak location within individuals across scanning sites has been estimated to vary by around 6 mm to 8 mm (Wurnig et al., 2013). Alternatively, peak activation locations may vary across participants, such that the population peak does not coincide with any single individual’s peak. In this case, bootstrap variability may reflect genuine variability in the location of peaks. In support of this, consider the variable success at localizing peaks within the UKB. The ten highest peaks were primarily located in more unimodal regions (Huntenburg et al., 2018), especially the visual cortex (see Table S3). Given that localization improves with larger effect sizes (Figure S7), smaller effect sizes could reflect increased peak spatial variability. Put another way, the existence of larger effect sizes in unimodal regions is consistent with less spatial variability in those regions, and conversely, greater variability in the transmodal areas. Nevertheless, as noted in subsection 3.2, this is likely an incomplete story, given that localizability of peaks within even unimodal regions varies strongly across tasks (Figure S6).

Finally, it is also possible that individual-level variability in native-space peak location can be reduced by improved normalization. The success of normalization methods that incorporate functional information (e.g., hyperalignment, MSMAll; Haxby et al., 2011) suggests that diffeomorphic transformations based solely on structural features result in misaligned functional features. Disentangling these cases is outside the scope of this report and would require repeated measurements within participants, across time and scan acquisition parameters.

Across most analyses, there was only a minimal difference between surface-based (MSMAll and SURFACE) and volumetric analyses. This may appear to conflict with the claim that surface-based analyses are preferable, especially regarding their spatial precision (e.g., Coalson et al., 2018). However, the analyses here were chosen because they represent ways to answer typical questions in neuroimaging (e.g., “which regions are differently activated by a given task?”, “is a characteristic predictable?”), answers that do not depend on high spatial resolution. For this reason, we are not making any claims about a lack of substantive advantages between the different data types. Some differences were observable in the reliability of peak detection. In particular, peak location was more variable in the surface-based analyses as compared to the volumetric analyses Figure S12, likely a product of the geodesic distance between voxels that are proximal to each other but also in separate anatomical regions (e.g., voxels on neighboring gyri). In some tasks, MSMAll resulted in peak locations with greater variability than SURFACE, although this does not necessarily reflect worse performance; if activation peaks vary substantially across participants, then higher variability may better reflect the ground truth.

Peaks capture only one feature of an activation map. Prior work has reported that most activation patterns in the Human Connectome Project are diffuse (i.e., spanning multiple anatomically defined regions) (Cremers et al., 2017). In this setting, the relevance of a single peak becomes less clear. When exploring the topography of the entire statistical map, we observed correlations between individual studies and their respective gold standards that ranged from strong to very strong ((0.5 - 1), see also Bossier et al., 2020; Sochat et al., 2015). For tasks that elicit substantial activation, maps constructed with only 20 participants showed correlations with the gold standard that exceeded 0.9. Thus, with respect to this global measure, even very small studies can provide information that is highly predictive of the broader population-level map.

Compare these results to those reported by the Neuroimaging Analysis and Replication Project (Botvinik-Nezer, 2020). In that project, teams of researchers analyzed the same set of data, each using its own idiosyncratic set of methods. A key finding of that project was that the analysis pipeline has a strong influence on binary activation maps (see also Bowring, Maumet, & Nichols, 2019), but a substantially weaker effect on the underlying unthresholded statistical maps. That is, when multiple analysis pipelines are applied to a common dataset, the unthresholded statistical maps are largely consistent with each other (M. Lindquist, 2020; Taylor et al., 2023). Similarly, we demonstrate that when a single analysis pipeline is applied to multiple repeated experiments, the statistical maps are consistent with each other.

Finally, the results using multivariate models were mixed. In general, tens of participants were sufficient for obtaining significant predictions on external training samples for some measures (Figure 7, Figure 8). However, the consistency of these predictions was poor (Figure 7b), as were the learned features (Figure 10b). This implies that the parameters learned by models remain unstable at these sample sizes. We speculate that this instability may relate to the high imbalance between the number of participants (in the tens) and the number of features (in the thousands). The models used regularization (ridge regression), but the particular regularization procedure aims to improve cross-validated performance, rather than feature stability. Data may provide several disjoint sets of features that can support equally good predictions (Adkinson et al., 2025). Therefore, without additional information, the regularization procedure may select different sets of features across studies. At no sampled level does the study-to-study reliability of features as measured by ICC exceed 0.4; that is, even 10,000 participants are too few participants to achieve better than “poor” reliability (Cicchetti, 1994).

### 4.1 Limitations

The analyses presented here focus on sample size, but there is likely a strong dependence of validity and reliability on the amount of high-quality data contributed by each participant. For example, scans half the duration of those analyzed here would yield noisier connectivity estimates (fewer time points, lower SNR), likely reducing out-of-sample modeling performance. In some study design configurations, the number of participants in a study is interchangeable with the time dedicated to scanning each participant (Ooi et al., 2025). Likewise, a larger sample built from participants who contribute low-quality data to a study (e.g., those who move excessively) may not yield a valuable dataset (e.g., because substantial sections of their scans must be scrubbed). Thus, when interpreting the specific quantities reported here, the amount of high-quality data provided by each participant must be taken into consideration. That is, our analyses should be interpreted with the caveat that they apply to datasets of comparable quality to the HCP-YA and UKB.

All modeling results relied on a single prediction method (ridge regression), and alternative methods will likely produce quantitatively different results. In particular, methods that achieve more stable features despite low sample sizes may be able to obtain higher feature stability (e.g., Du et al., 2020).

Several modeling results showed substantial study-to-study reliability differences between UKB and HCP-YA volumetric datasets (Figure 7b, Figure 8, Figure 10a, Figure 10b). There are substantive reasons why the UKB may be different than the HCP-YA counterpart (task: emotion); the population was much more diverse, average motion differed, and the size of the dataset means that there is less opportunity for QC of individual scans. Each of these differences may conspire to lower validity and reliability. However, we also highlight that the subsampling method adopted here and in other studies is susceptible to bias, such that a smaller population (e.g., 400 HCP participants vs 40,000 UKB participants) leads to higher estimates of validity and reliability. That is, some of the differences between the UKB and the HCP-YA emotion task are likely driven by methodological issues. Consider the reliability of model predictions (Figure 7b). When samples are drawn from a relatively small population (e.g., the HCP-YA dataset), it is likely that the samples will contain the same participants. With overlapping sets of training data, the resulting models will produce predictions that are more similar than they would have been if the models had been trained on disjoint sets of data. We elaborate on the issue in subsection 5.3, and it will be explored in future work. Here, we provide the caveat that the absolute values for measures of validity and reliability may be biased upwards. In analyses where there is a relatively small difference between the UKB and HCP-YA results, the bias due to this methodological issue is not expected to be substantial. Regardless, that caveat does not change the main conclusion regarding the modeling results, which is that even 100 participants is likely too few for most modeling analyses (any analysis beyond the question “is characteristic Y predictable from dataset X?”), especially when study-to-study reliability is vital for conclusions.

### 4.2 Recommendations

First, we continue to remind neuroimaging researchers that data ought to be made publicly available. There have been calls for open sharing for over a decade. Although tools have been developed to work around the lack of readily available raw images or statistical maps (e.g., neurosynth.org), and community-driven efforts demonstrate the feasibility of decentralized data sharing (e.g., the FCON 1000 project), numerous resources now exist that obviate these workarounds. In the US alone, these resources include OpenNeuro, the National Institute of Mental Health Data Archive, and NeuroVault (Gorgolewski et al., 2015). The availability of rich and varied raw data substantially increases the value of small studies, especially when analyses are exploratory or aim to probe subtle effects.

Second, for specific well-circumscribed aims, we recommend against overemphasizing lack of reproducibility; datasets with tens of participants, which are typical in the neuroimaging literature (Poldrack et al., 2017), can be of high quality and value. For well-studied tasks that produce large effect sizes (e.g., the language, motor, social, and working memory tasks of the HCP dataset), 40 participants provide high power to detect regional activation. However, we emphasize that this recommendation applies to datasets of comparable quality to the HCP, with tasks that are known to produce at least medium effect sizes, and with limited room for exploratory analyses. In tasks with novel effects or unknown effect sizes, tens of participants are likely too few to warrant confidence in a new effect, or in which regions are most activated by the task. Even so, the required sample sizes may not be in the hundreds, considering that around 80 participants were enough to reliably activate the targeted regions even in the most challenging tasks (gambling and relational).

Third, there is a need for further research on quantifying confidence in peak location. The reported distributions of peak distances provide heuristics for assigning confidence to locations reported in individual studies. Still, these heuristics imply a general uncertainty (e.g., with 40 participants, any voxel within 10 mm of a reported peak is a likely location for the true peak). Typical cluster analyses discard substantial information about the location of activation, given that the significance of a cluster only implies that there is an activation in some voxel within a cluster (Woo et al., 2014). This can lead to situations where larger study populations increase the power to detect activation within each voxel, thereby increasing the size of clusters and hindering the determination of which voxels are active (Rosenblatt et al., 2018). Worse, the question of whether there is a voxel above 0 is qualitatively different than the question implied by an assessment of peak location, which is whether the activation in a voxel is significantly higher than the activation of its neighbors. Advances have been made in exploring confidence in effect size maps (Bowring, Telschow, et al., 2019; Bowring et al., 2021), but these methods are not yet commonly used, and so it is not yet clear how they perform in a wide range of datasets.

Finally, we advise against using predictive models that have been trained on data from tens of participants in any applied or clinical setting. This recommendation derives from the study-to-study comparisons of modeling. Across training datasets, models trained on small sample sizes make predictions that have poor consistency (Figure 7b), potentially resulting in clinical decisions that would be highly dependent on particular training samples. Moreover, not only are the predictions unstable, but the features themselves are also unreliable. Low feature reliability means that, without external information, feature importance within a model provides little justification for the relevance of that feature to the predicted entity.

## Data and Code Availability

Code to reproduce analyses is available on GitHub: https://github.com/psadil/maps-2-models. Analyses relied on open data provided by the Human Connectome Project, which can be downloaded from the HCP website https://humanconnectome.org/study/hcp-young-adult/document/500-subjects-data-release, and on the UK Biobank.

## Author Contributions

**Patrick Sadil**: Conceptualization, Methodology, Software, Validation, Formal Analysis, Investigation, Resources, Data Curation, Writing - Original Draft, Writing - Review & Editing, Visualization. **Martin A. Lindquist**: Conceptualization, Methodology, Validation, Formal Analysis, Resources, Writing - Original Draft, Writing - Review & Editing, Supervision, Project Administration, Funding Acquisition.

## Ethics Statement

Informed consent was obtained from all Human Connectome Project participants.

## Acknowledgements

Data were provided by the Human Connectome Project, WU-Minn Consortium (Principal Investigators: David Van Essen and Kamil Ugurbil; 1U54MH091657), funded by the 16 NIH Institutes and Centers that support the NIH Blueprint for Neuroscience Research, and by the McDonnell Center for Systems Neuroscience at Washington University.

This research has been conducted using data from UK Biobank, a major biomedical database (Project ID: 33278). We are grateful to UK Biobank and the UK Biobank participants for making the resource data possible, and to the data processing team at Oxford University for sharing the processed data. The UK Biobank imaging project is funded by the Medical Research Council and the Wellcome Trust.

## 5 Supplementary Material

### 5.1 Excluded Participants

The following 93 participants were excluded from analyses due to either QC issues or the presence of a twin in the included sample: 110613, 113417, 113821, 120010, 121719, 130518, 139637, 143830, 146836, 168139, 175035, 176239, 185038, 189652, 199958, 201515, 202820, 385046, 401422, 415837, 433839, 462139, 465852, 469961, 644246, 656657, 688569, 723141, 767464, 872764, 943862, 965367, 969476, 987983, 994273, 433839, 103010, 113417, 116423, 120010, 121719, 127226, 130114, 143830, 169040, 185038, 189652, 202820, 204218, 329844, 385046, 401422, 462139, 469961, 644246, 688569, 723141, 908860, 943862, 969476, 971160, 196952, 748662, 809252, 144428, 186545, 192237, 223929, 320826, 644044, 822244, 870861, 947668, 102614, 111009, 111514, 115017, 121416, 130821, 138332, 179952, 299760, 300618, 392750, 406432, 429040, 633847, 662551, 679770, 688569, 693461, 815247, 142626

### 5.2 Tasks

#### 5.2.1 HCP

The following task descriptions are paraphrased from the details provided by Barch et al. (2013).

##### Working Memory

In the working memory runs, participants (494) completed N-back tasks (Drobyshevsky et al., 2006). Each of the two runs contains eight stimulus blocks consisting of ten trials (2 s stimulus presentation, 500 ms ITI) and four fixation blocks (15 s each). Within each run, four different stimulus types (faces, places, tools, and body parts) were presented in separate blocks. Task type was indicated at the start of each block with a 2.5 s cue. Each block contains two targets and 2–3 non-target lures. Half of the blocks for each stimulus type used a 2-back task, and the other half a 0-back task. Total scan time was 5:01 (405 frames).

The design matrix included eight task-related predictors, one for each stimulus type in each of the N-back conditions. Each predictor covered the period from the onset of the cue to the offset of the final trial. Our analyses considered a contrast selecting for the 2-back parameter, the 0-back parameter, and a comparison of the 2-back minus 0-back parameters.

##### Motor

In each of two runs, participants (492) were asked to move different body parts (Buckner et al., 2011; Thomas Yeo et al., 2011). The body part to move was indicated by visual cues presented at the start of each block (one 3 s cue for each 12 s block). Cues indicated that participants should either tap their left or right fingers, squeeze their left or right toes, or move their tongue (only one type of motion was asked for in each block). Each run contained two blocks of tongue movements, two blocks of each hand movement, two blocks of each foot movement, and three additional blocks of fixation (each 15 s). Total scan time was 3:34 (284 frames).

The design matrix included five task-related predictors, each covering the duration of the 10 movement trials (12 s). The cue was modeled separately. Our analyses included a contrast selecting for the cue, one selecting for the average motion, and a comparison of the average motion minus the baseline.

##### Gambling

In this task, participants (494) were asked to guess the number on a hidden card to win or lose money (Delgado et al., 2000). They were informed that the number ranged from 1 to 9, and that they should guess whether the hidden number was greater or less than 5. Guesses were realized by pressing one of two buttons. The task was presented in blocks of eight trials that comprised either mostly rewards (6 reward trials interleaved with either one neutral and one loss trial, two neutral trials, or two loss trials) or mostly losses (6 loss trials interleaved with either one neutral and one reward trial, two neutral trials, or two reward trials). After responding, participants were given feedback. On reward trials, the feedback was a green up arrow with $1, on loss trials, it was a red down arrow with $0.50, and on neutral trials, it was the number 5 with a gray, double-headed arrow. In each of the two runs, there were two mostly reward and two mostly loss blocks, interleaved with four fixation blocks (each 15 s). Total scan time was 3:12 (253 frames).

The design matrix included two task-related predictors that modeled the mostly reward and mostly punishment blocks, each covering the duration of 8 trials (28 s). Our analyses relied on contrasts for the reward parameter, the punish parameter, and the reward minus punish parameters.

##### Language

In each run of this task, participants (483) listened to eight blocks of stimuli, four of which consisted of short stories and four of which consisted of arithmetic (Binder et al., 2011). Blocks averaged 40 s and the two stimulus types were interleaved. After each block, participants were presented with a two-alternative forced-choice question that either asked about the story or the result of the arithmetic. Total scan time was 3:57 (316 frames).

The design matrix included two predictors that corresponded to the two types of stimuli. Our analyses relied on contrasts for the story parameter, the math parameter, and the difference between math and story.

##### Relational

In this task, participants (481) were presented with sets of stimuli and asked to discern whether they matched or differed according to pre-specified rules (R. Smith et al., 2007). Stimuli varied by shape and texture. In a relational condition, two pairs of stimuli were presented. One pair serves as a reference, and the stimuli in that pair differ along one of the two dimensions. Participants first identified the mismatching dimension and then determined whether the other pair of stimuli differed along the same dimension. In a matching condition, a single pair of object stimuli was presented along with a third object and a word. The word identified one of the two features, and participants had to determine whether the identified feature in the third stimulus matched the feature value for either of the paired stimuli. Relational stimuli were presented for 3500 ms (ITI: 500 ms) and matching stimuli for 2800 ms (ITI: 400 ms). Stimuli were presented in three blocks that each contained five trials, and the stimulus blocks were interspersed with three fixation blocks (16 s). Total scan time was 3:57 (316 frames).

The design matrix for this task included two predictors, one for each of the two conditions (“match” and “relation”). Our analyses relied on contrasts for the matching minus relation parameters.

##### Social

The social task used stimuli derived from Castelli et al. (2013) and Wheatley et al. (2007). The stimuli were videos of shapes that moved either randomly or with a set of specified interactions (20 s per video). After each video, participants (486) selected one of three responses indicating whether they observed the objects interacting. There were five blocks of trials per run (conditions balanced across the two runs). Total scan time was 3:27 (274 frames).

The design matrix for this task contained two predictors, one for each of the “theory of mind” and “random” conditions. Our analyses used contrasts that selected the subtraction of the theory of mind from the random parameter.

##### Emotion

In this task, participants (374) were presented with blocks of trials that either asked them to decide which of two faces shown at the bottom of the screen matched the face at the top, or which of two shapes presented at the bottom matched the shape at the top. The faces had either angry or fearful expressions. Trials are presented in blocks of 6 trials (2 s stimulus presentation, 1 s ITI) of the same task (“face” or “shape”). Each block was preceded by a 3 s task cue (“shape” or “face”). Each of the two runs includes three face blocks and three shape blocks. Total scan time was 2:16 (176 frames).

Two different task-related predictors were included in the design matrix, one corresponding to emotional faces and the other to the shape control condition. Each predictor covered a 21 s duration composed of a cue and six trials. A linear COPE comparing emotional faces vs shapes was used for further analysis.

#### 5.2.2 UKB

The following task descriptions are paraphrased from the details provided by Alfaro-Almagro et al. (2018).

##### Emotion

The single UKB task is very similar to the Emotion task in the HCP. Both block types (faces and shapes) were each presented for five, 21 s blocks. Total scan time was 4:13 (332 frames). Analyses focused on the Faces-Shapes contrast.

### 5.3 Impact of Population Size on Simulations

As noted in the main text, the absolute values reported for reliability may exhibit a bias. Many of the analyses relied on a subset of the HCP-YA dataset with fewer than 400 participants. From this population, studies were generated by repeatedly drawing individuals, and the main results consisted of summaries derived from the samples. Here, we outline the source of the bias.

The key issue relates to sample overlap. When generating studies that are close to the full population size, the participant composition of each sample will have a large overlap. For example, with a population of 100 and a sample of 90, bootstrap studies will share substantial participant overlap. By contrast, with a population of 1,000 and the same sample size (90), overlap is much smaller, and many participants will appear in relatively few studies. Greater overlap artificially inflates between-study similarity, biasing reliability estimates upward.

To assess the impact of this, a simulation study was conducted. We considered the effect of dataset size on the reliability of voxel-wise measurements of effect size. We generated 100 “full” datasets (each participant contributing 100 voxels, with effect sizes generated from a normal distribution with mean zero and standard deviation 0.1), each with *M* participants contributing *P* = 100 observations. For each full dataset, *B* = 100 bootstrap samples were generated, each containing *N* = 16 participants. Each bootstrap sample was then summarized (a voxelwise average across the *N* participants), and the summaries were compared to each other by calculating an intraclass correlation. The correlations, shown on the y-axis, were then summarized across each of the complete datasets (mean and two standard errors). As the population size increases, the estimated reliability decreases (Figure S1). This is consistent with the interpretation that overlap in small finite populations inflates reliability.

**Figure S1:**
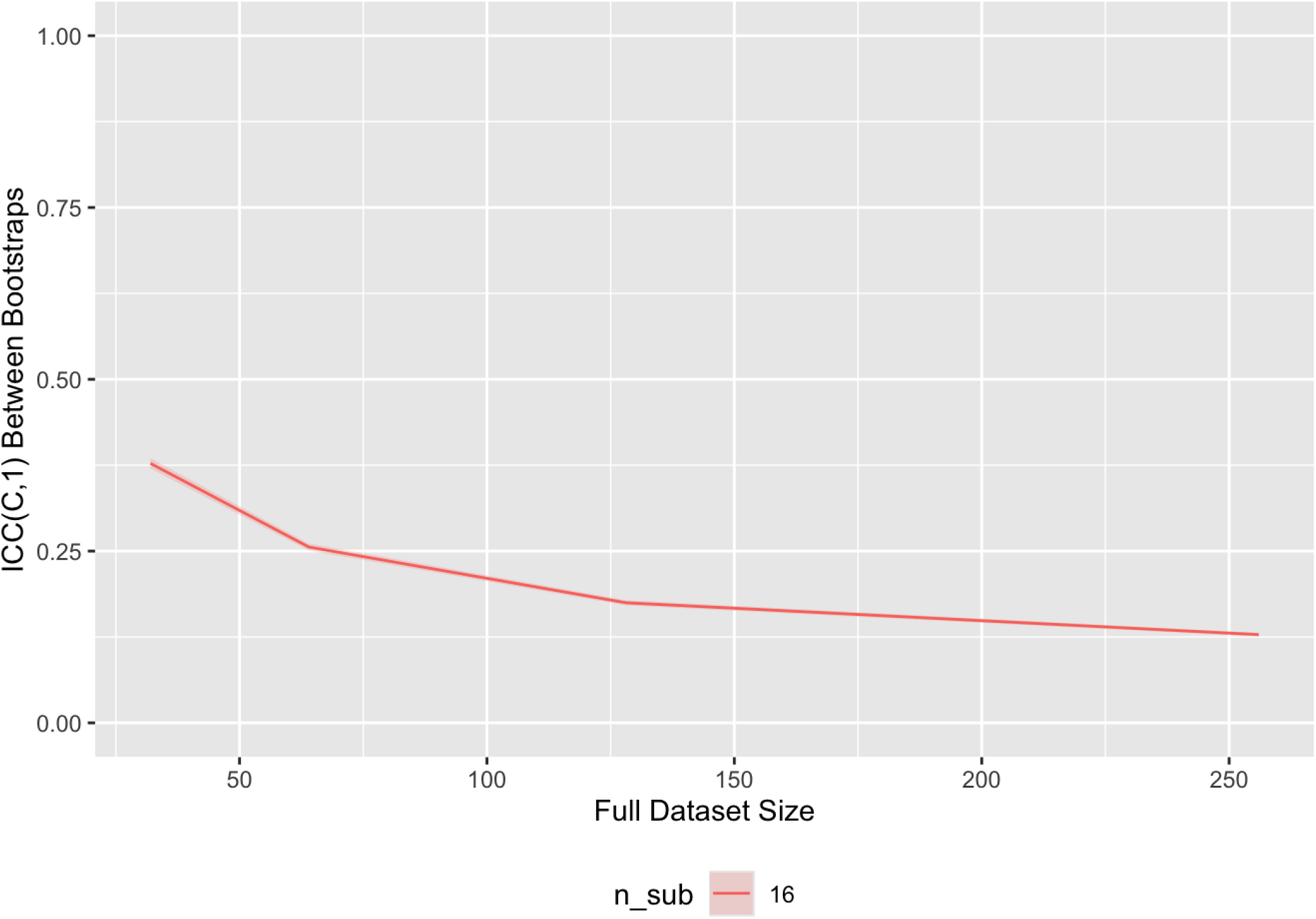
Estimated reliability of voxelwise effect sizes depends on sample size. The Intraclass Correlation Coefficient (ICC) was calculated as described in the main text.

### 5.4 Supplementary Figures

**Figure S2:**
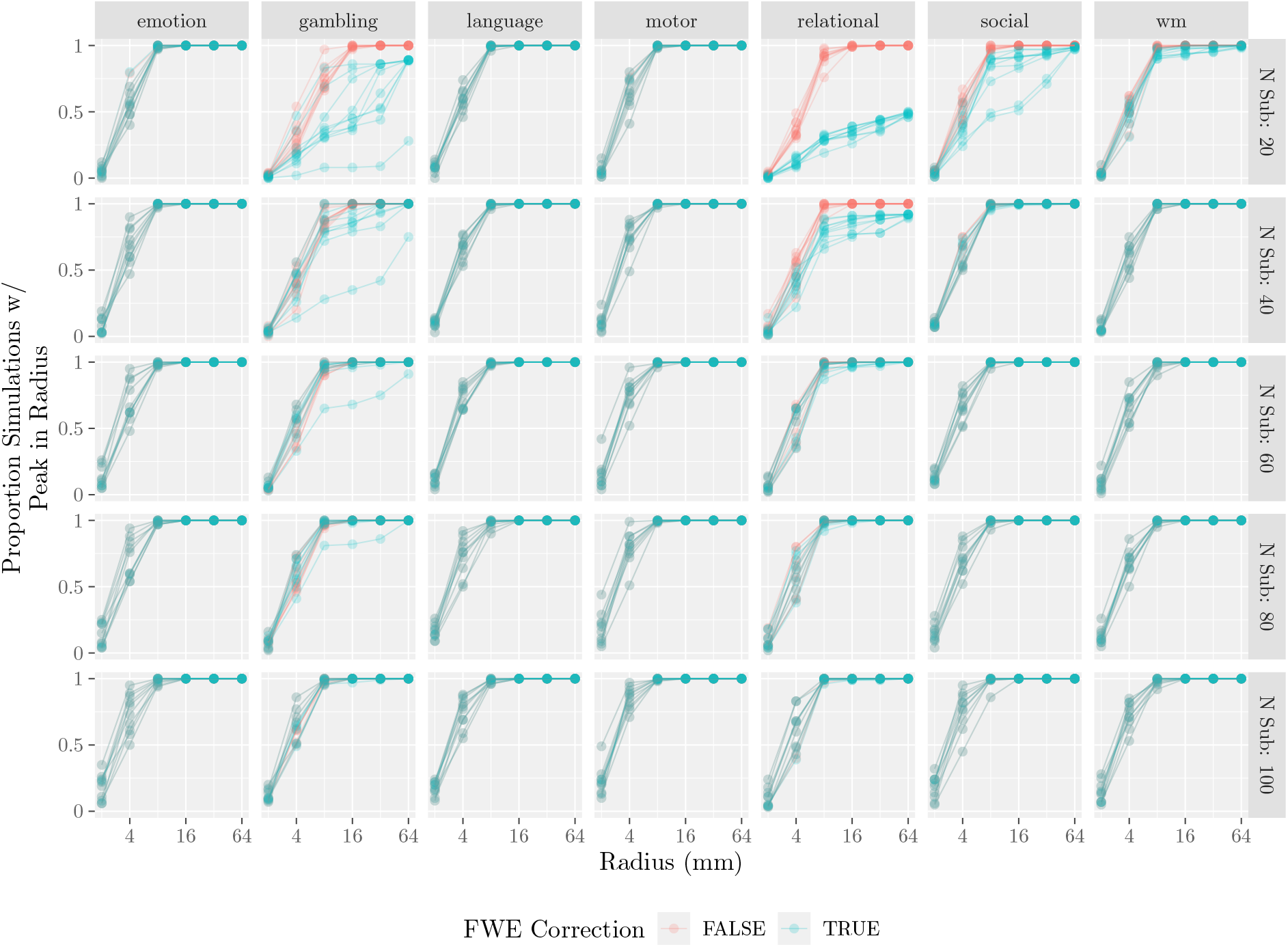
Volumetric peak localization with and without family-wise error rate correction using probabilistic threshold-free cluster enhancement (Spisák et al., 2019). Data plotted as in Figure 4. There is a clear effect of thresholding with family-wise error correction when sample sizes are 40 or 20, but at larger sample sizes, the effect is diminished. With more than 40 participants, there is only a minimal change in peak recovery for most of the largest peaks.

**Figure S3:**
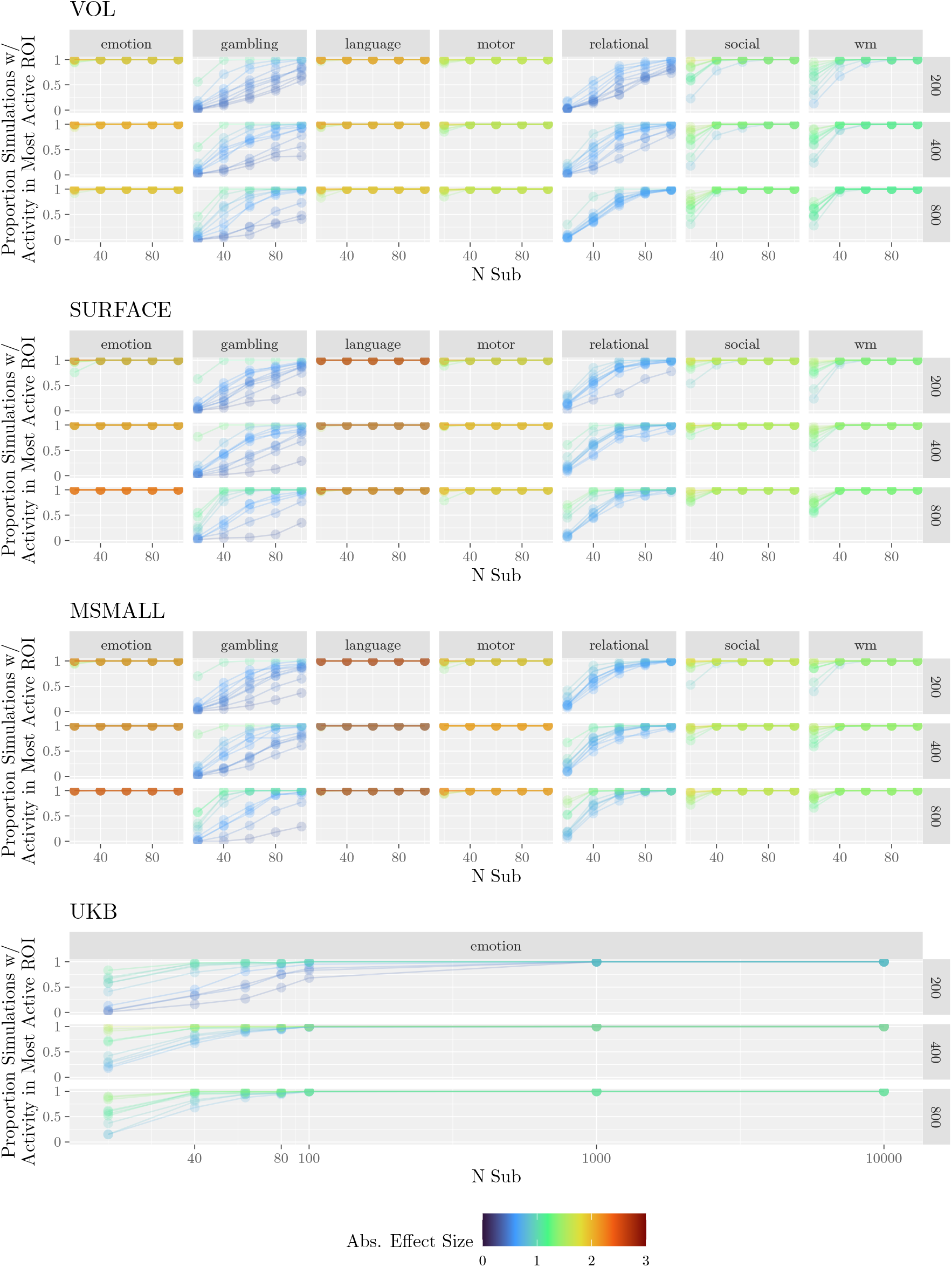
Recovery of gold standard across parcellations. Rows indicate the number of parcels within the atlas. Compare with Figure 2a.

**Figure S4:**
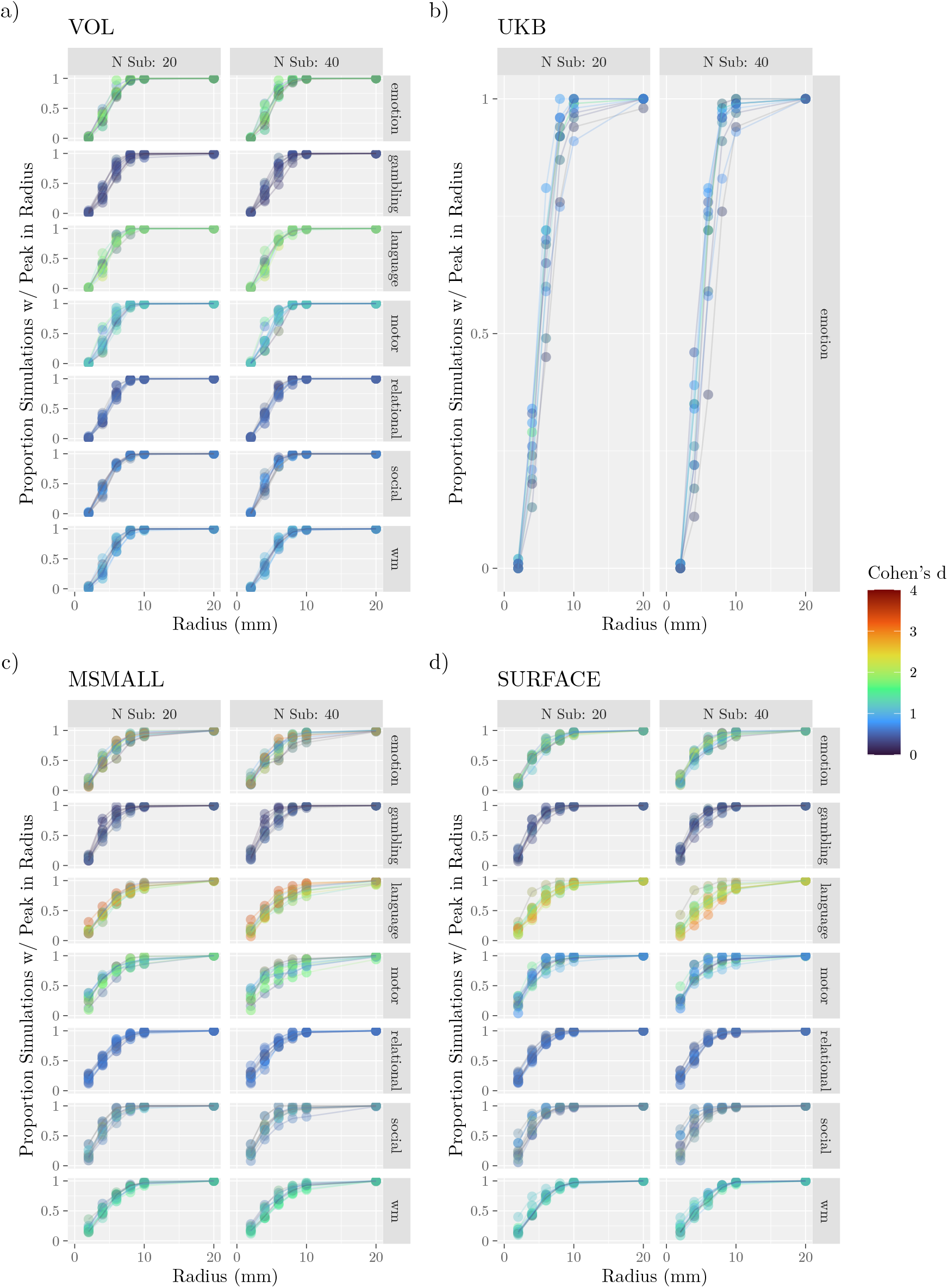
Figure plotted as in Figure 4, but peaks are defined within unthresholded maps.

**Figure S5:**
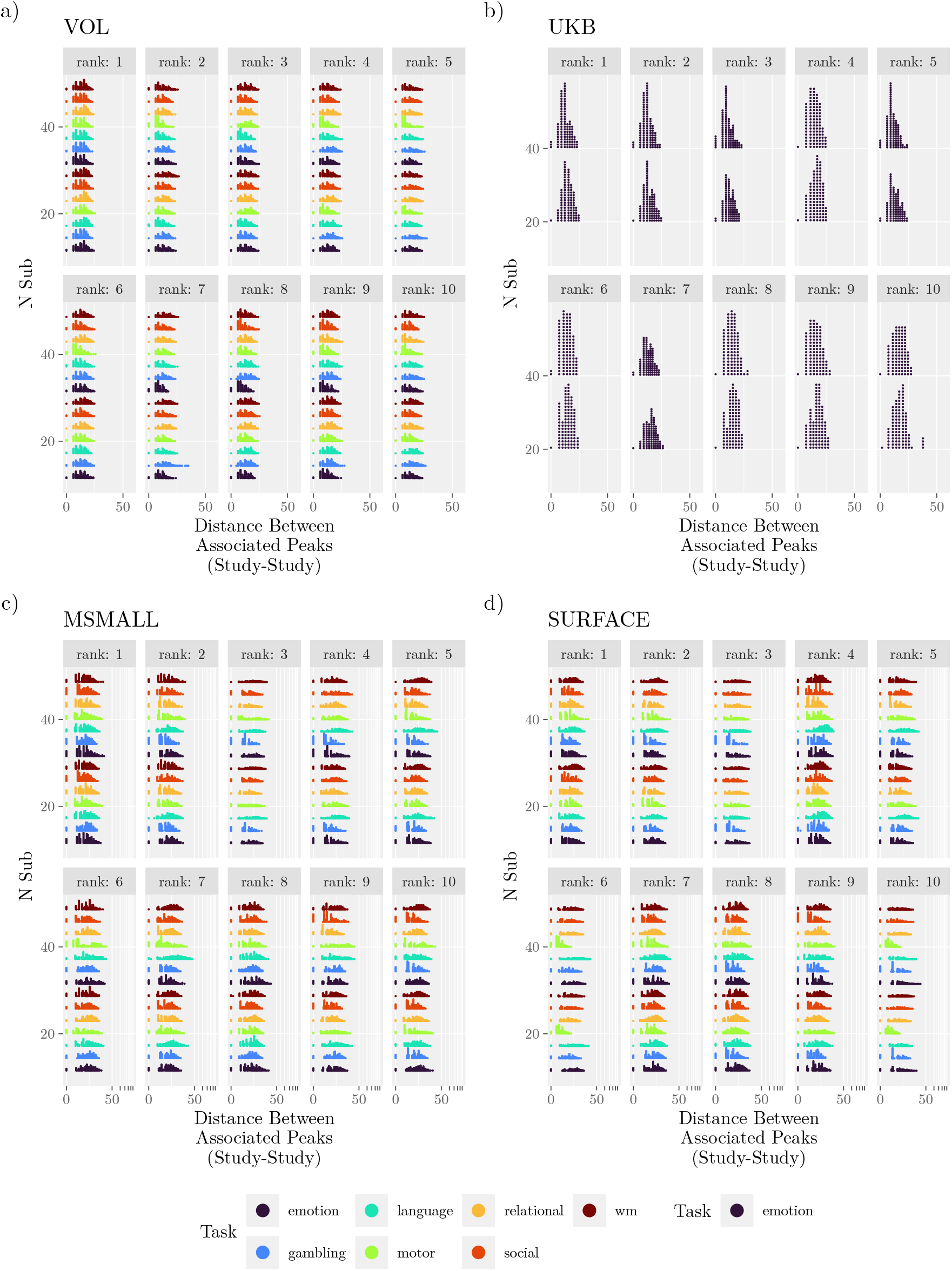
Figure plotted as in Figure 5, but peaks are defined within unthresholded maps.

**Figure S6:**
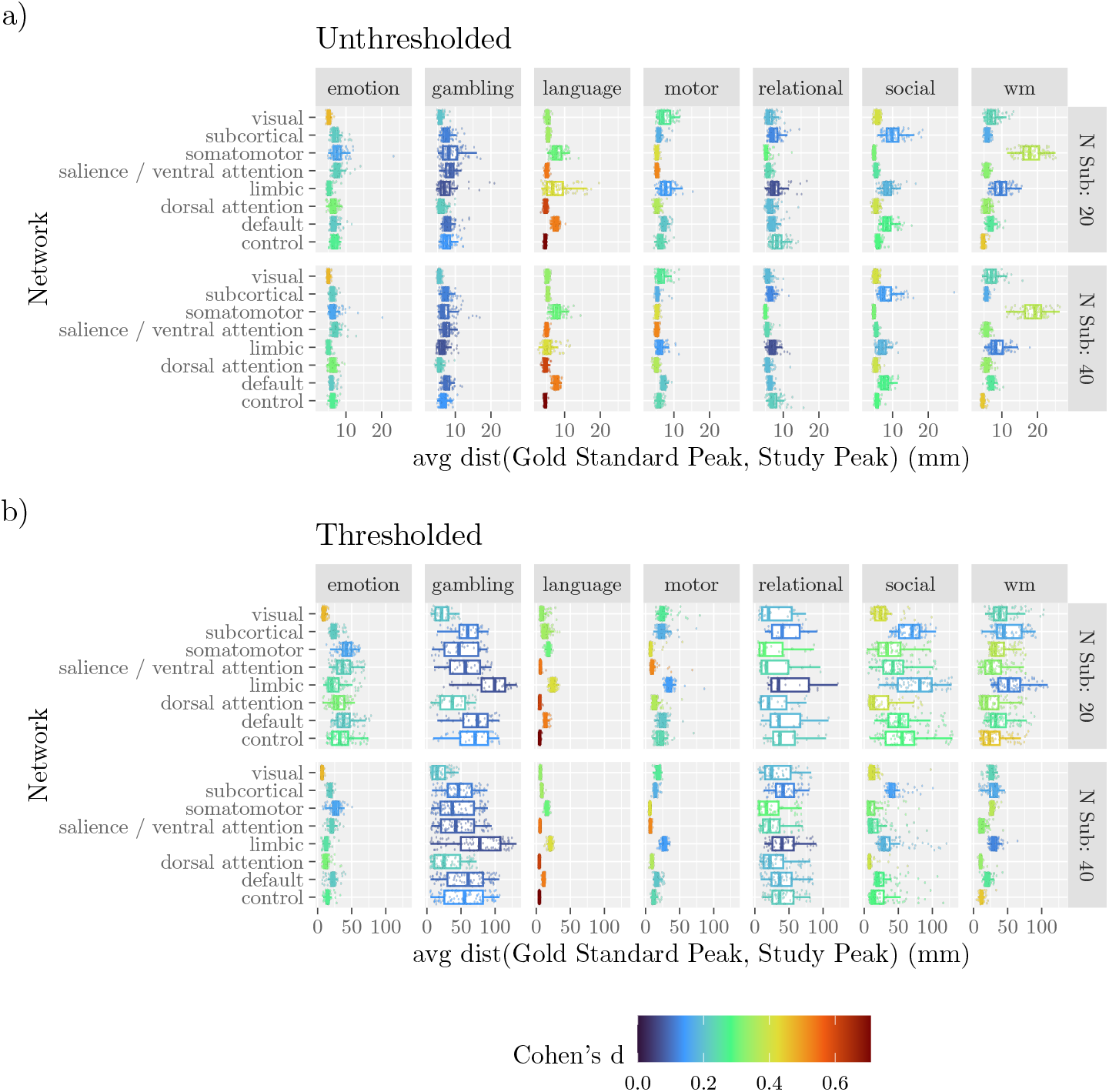
Distribution of average distances within Yeo7 networks. Studies are organized by sample size (rows) and task (columns). Peaks were selected from each gold standard map and labeled according to the Yeo7 networks (VOL only). Across all peaks within a network, the distance to the nearest study peak was calculated, and then these distances were averaged by study, network, task, and sample size. Points mark the distance to the nearest peak within each generated study, where the study maps were either unthresholded **(a)** or thresholded with family-wise error rate corrected *p <* 0.05 **(b)**. Colors indicate the average effect size of peak voxels in the gold standard for the given network.

**Figure S7:**
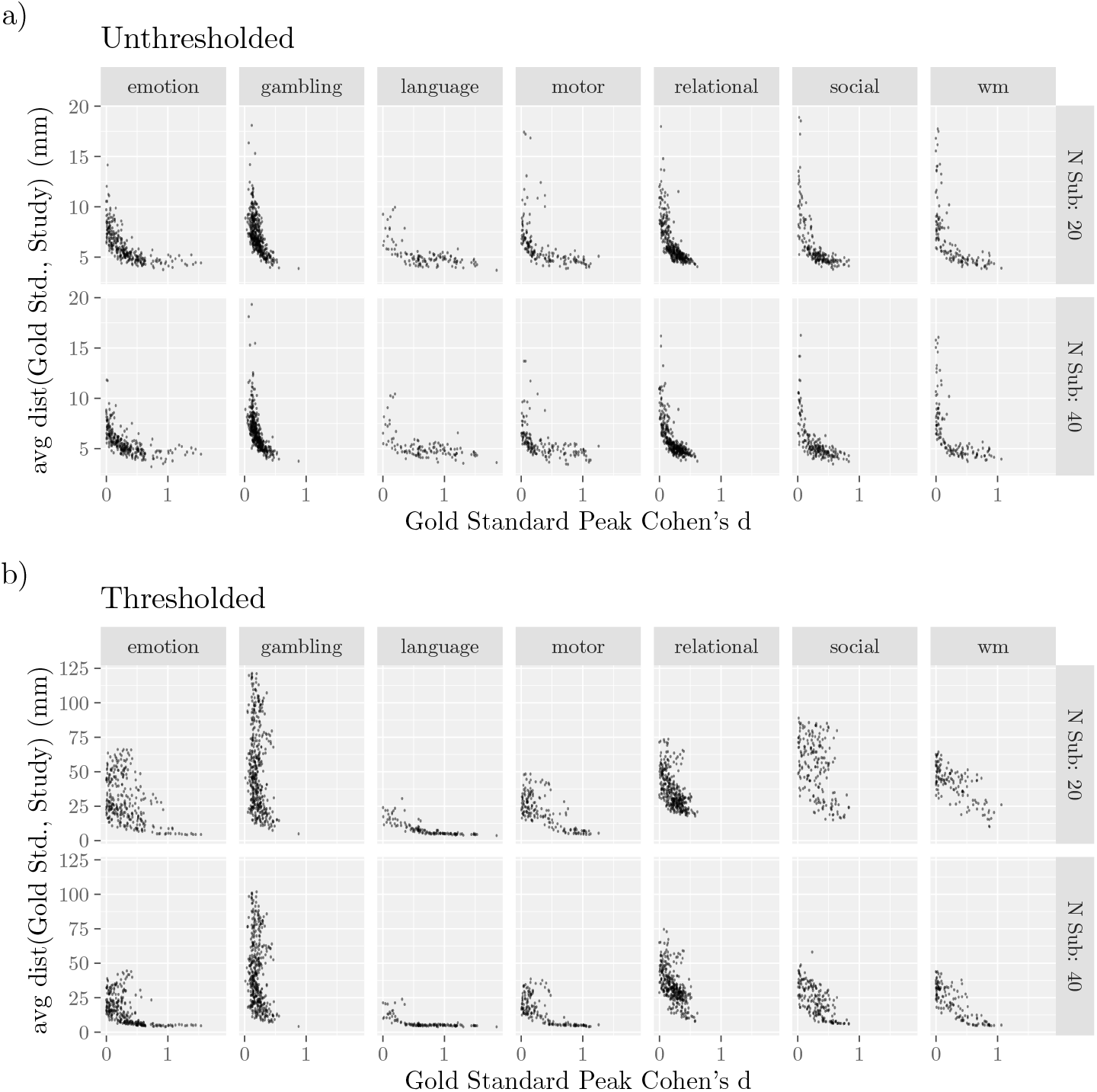
Average Distance from Gold Standard Peak by Effect Size. Individual points correspond to peaks in the gold standard. Averages were taken across studies and grouped by sample size. Note that only the VOL analyses are shown. Peaks are from maps that were either left unthresholded **(a)** or thresholded with family-wise error rate corrected *p <* 0.05 **(b)**.

**Figure S8:**
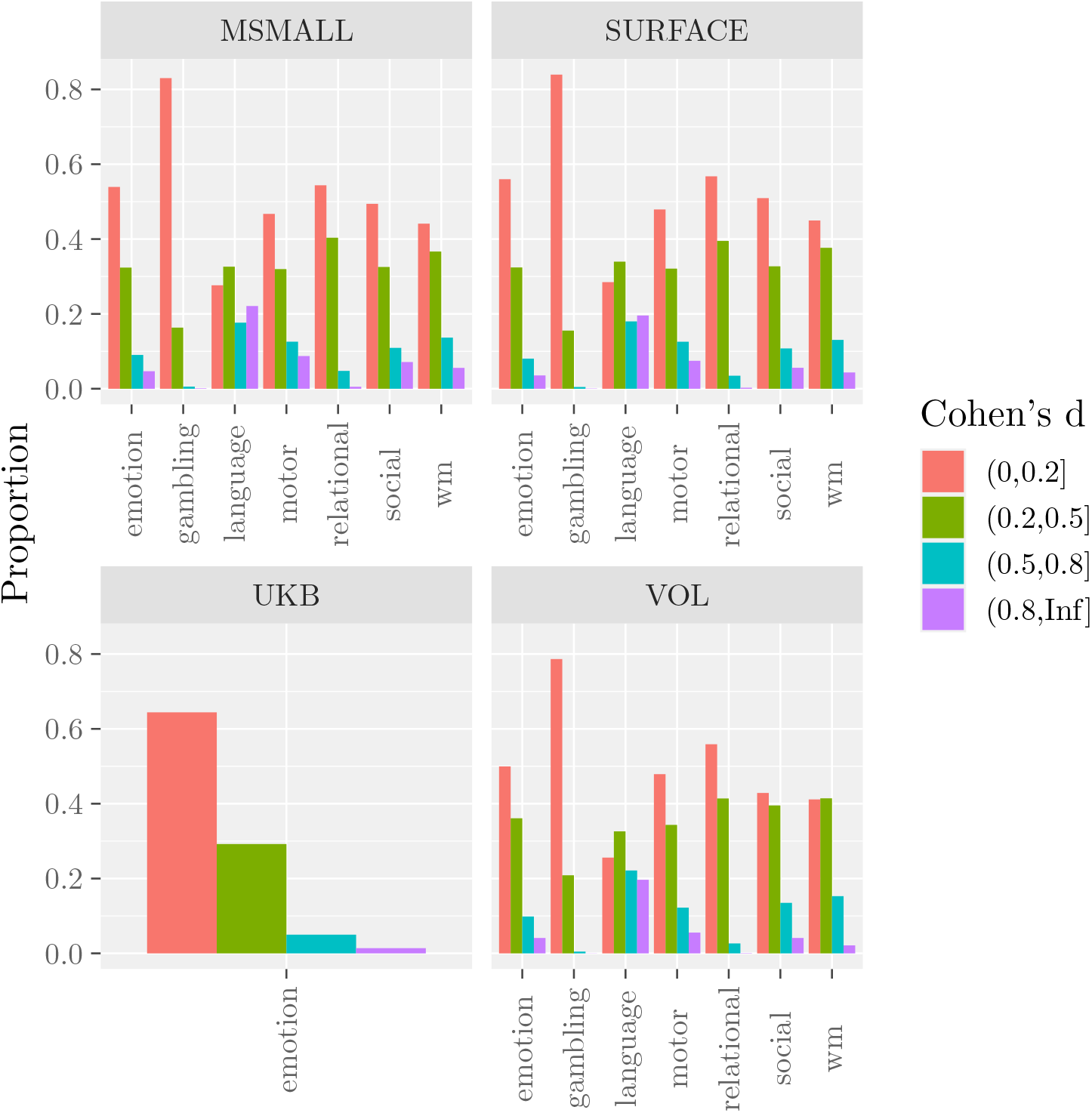
Voxel-wise effect sizes produced by each task. Categories were defined as specified in the Methods.

**Figure S9:**
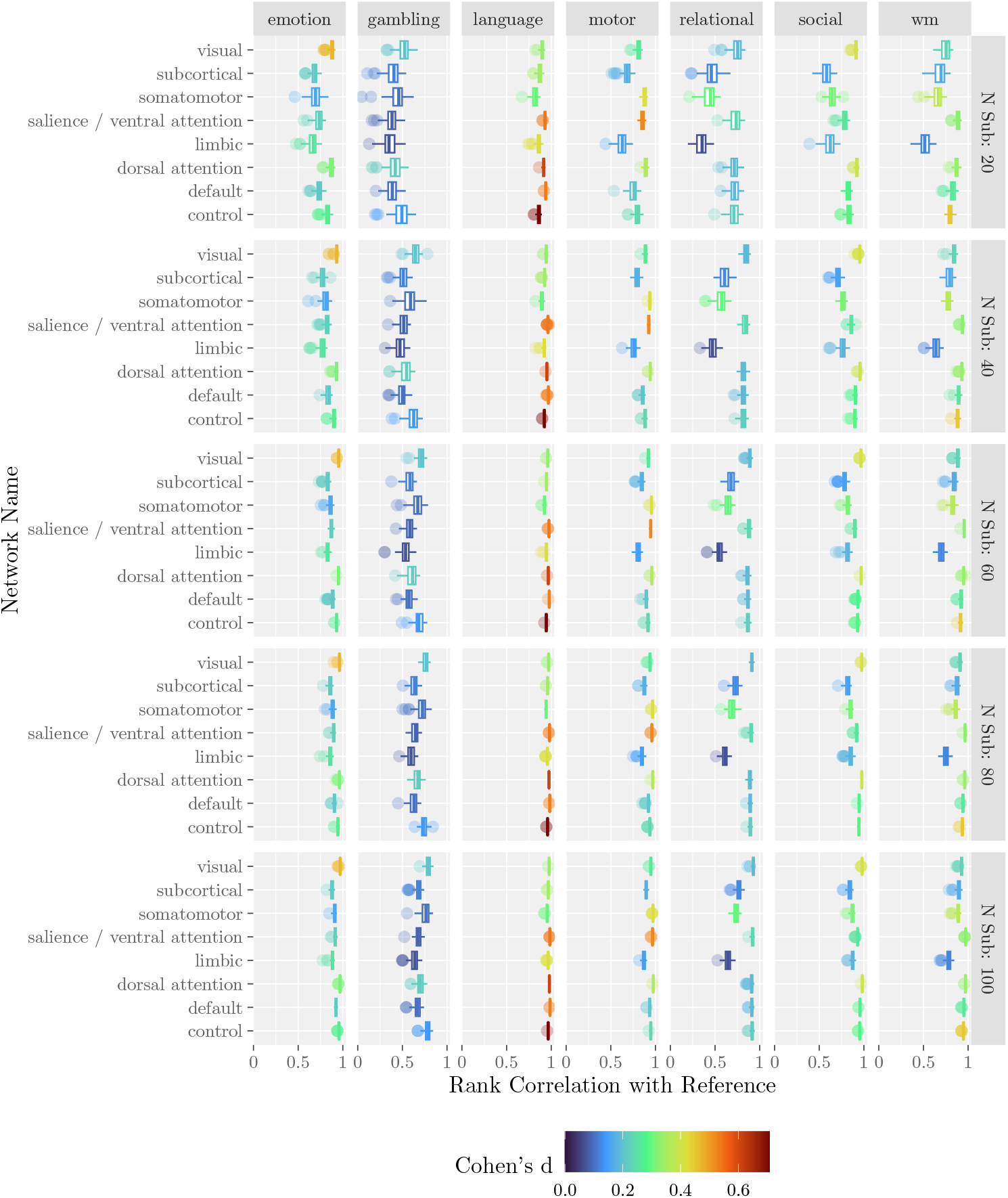
Recovery of gold standard by Yeo7 networks. Colors indicate the average effect size of voxels assigned to the network within the gold standard. Points correspond to sampled studies.

**Figure S10:**
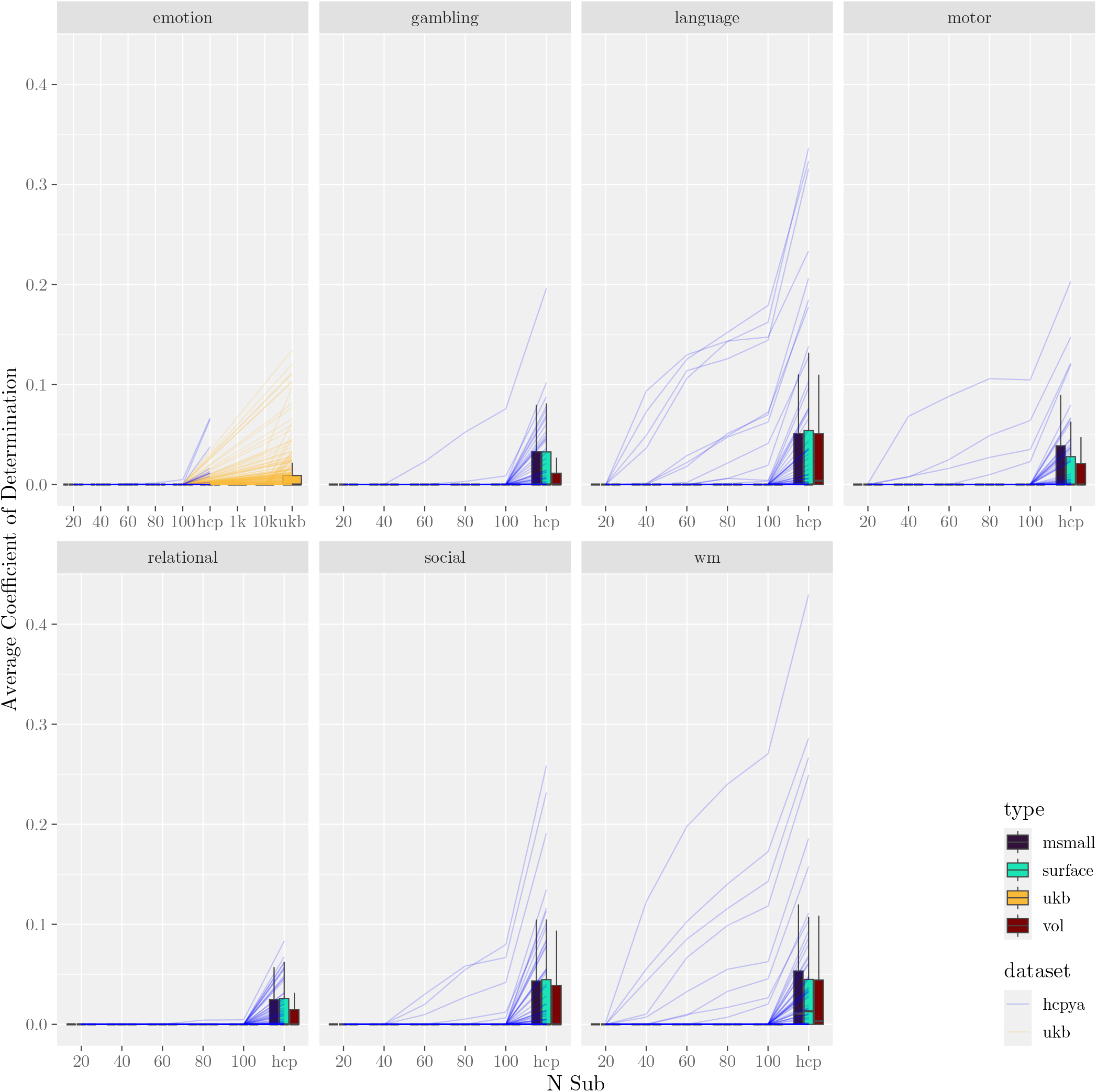
Multivariate model performance. Data plotted as in Figure 7a, but model performance is measured with *R*^2^. Note that values of *R*^2^ *<* 0 were set to 0.

**Figure S11:**
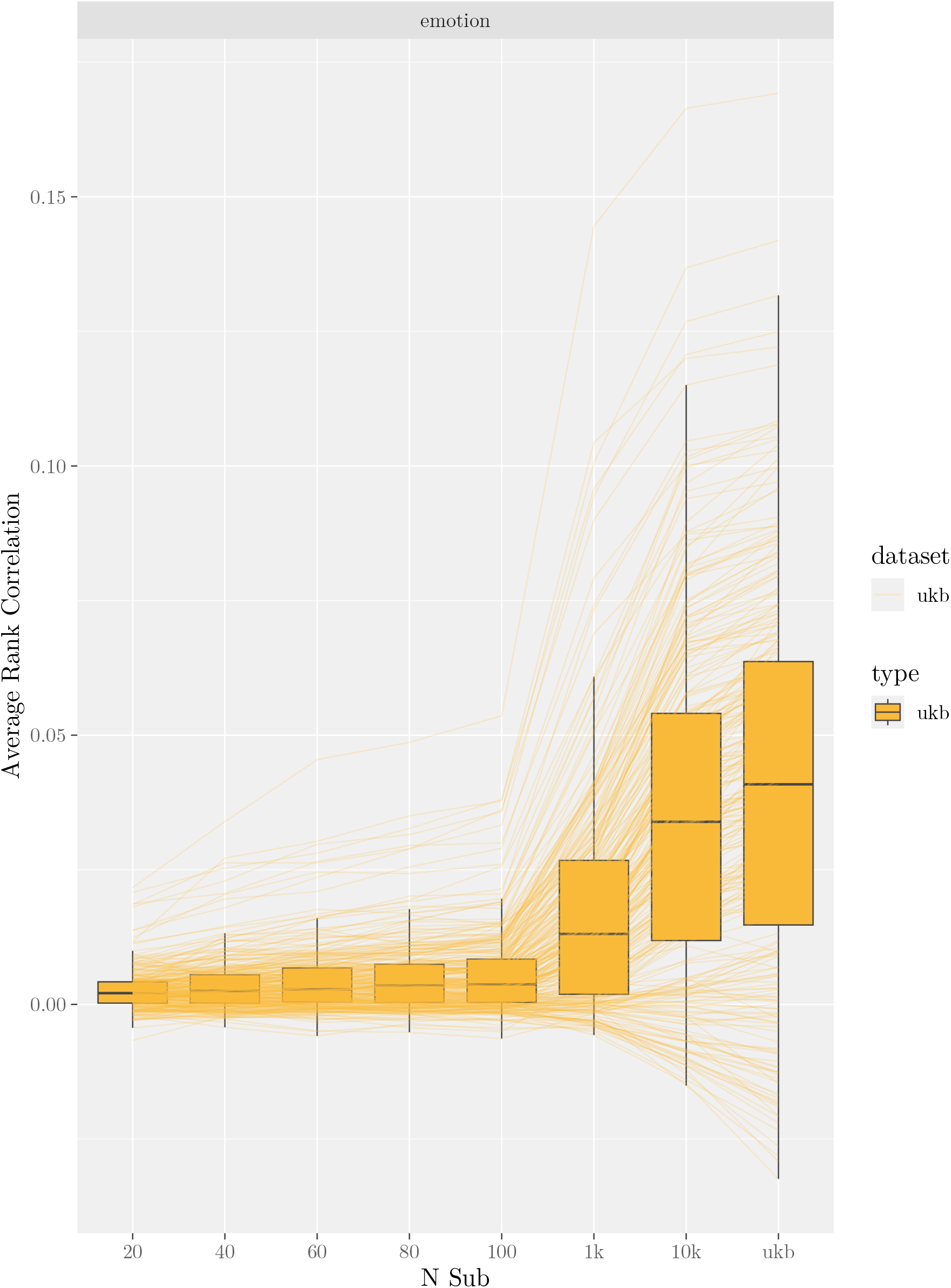
Model Performance with Alternative Features (Effect Size in 10 Highest ROI). Compare with Figure 7a.

**Figure S12:**
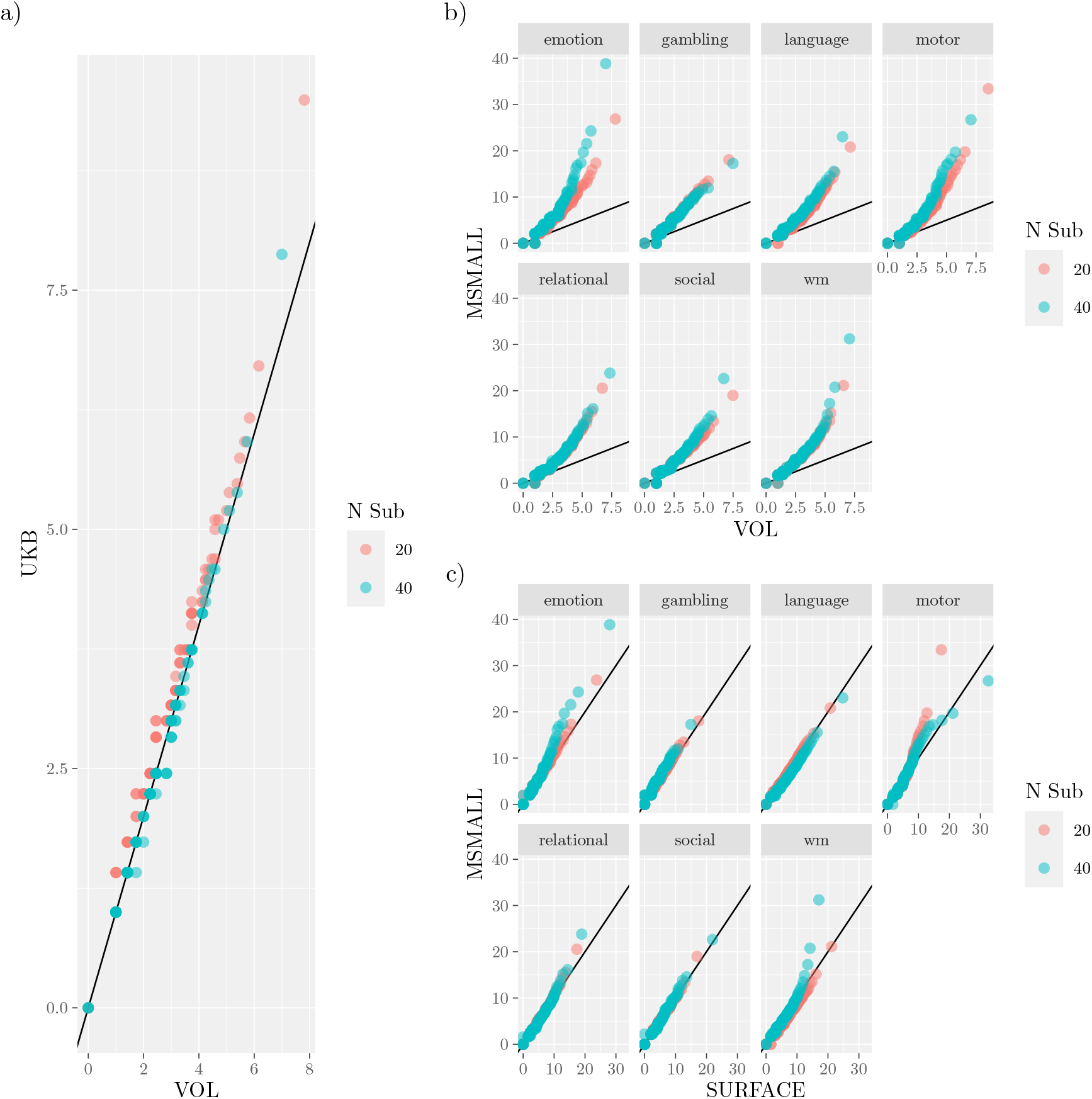
Differences in Peak Reliability Across Types. For each type, the distribution of distances for peaks associated with the largest peak activation (i.e., 4950 distances) was used to calculate an empirical cumulative density function. That function was then used to estimate the distribution percentiles, and these percentiles were plotted against one another. Solid lines mark the line of equality, so points above (below) the line indicate that the variable on the y-axis has greater (lower) reliability. a) VOL vs UKB. b) VOL vs MSMAll. c) Surface vs MSMAll.

**Table S1:**
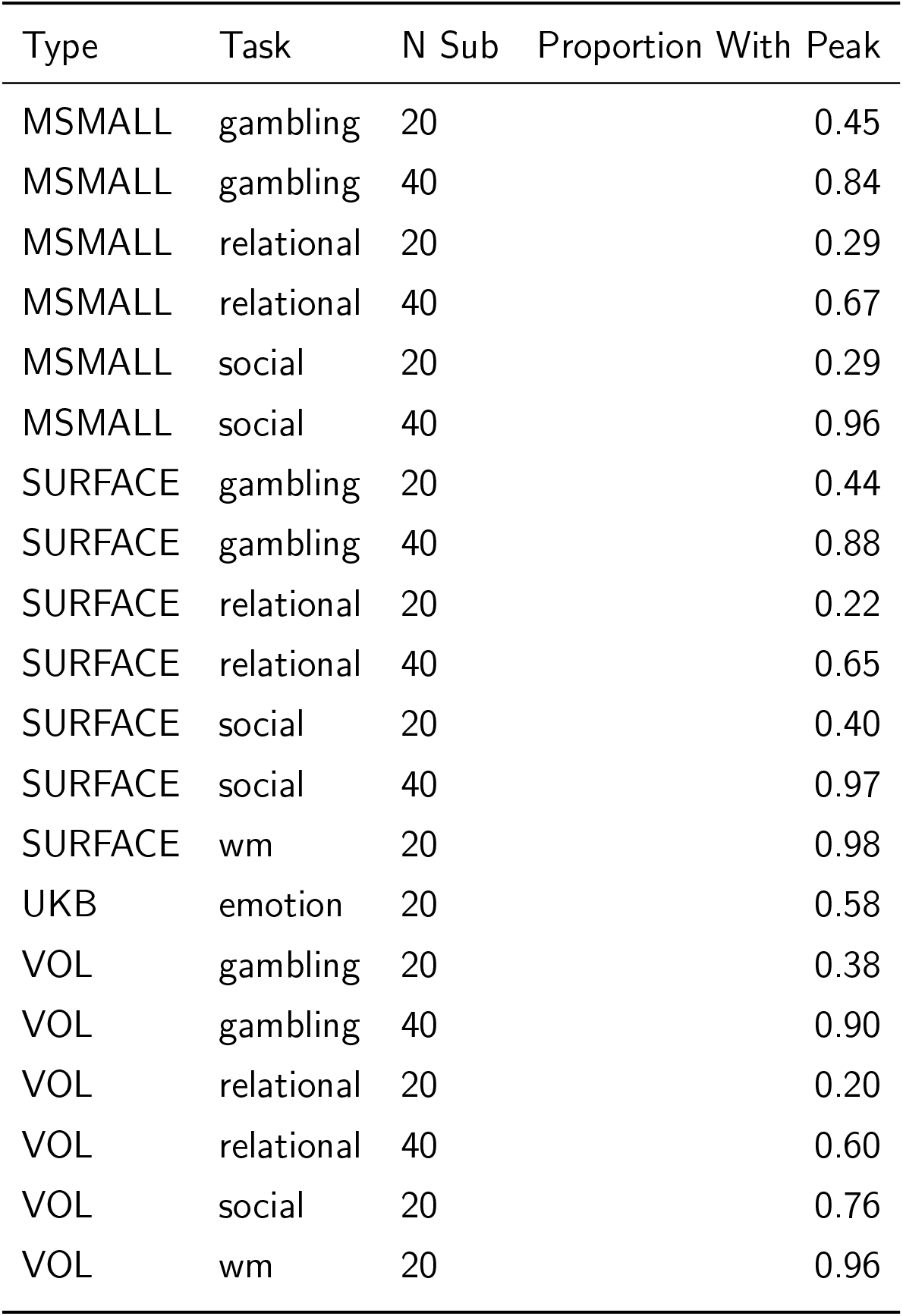
Proportion of Studies with Peaks Above Threshold. Entries with Proportions equal to 1 are excluded.

